# Barley resistance and susceptibility to fungal cell entry involve the interplay of ROP signaling with phosphatidylinositol-monophosphates

**DOI:** 10.1101/2024.11.19.624325

**Authors:** Lukas Sebastian Weiß, Mariem Bradai, Christoph Bartram, Mareike Heilmann, Julia Mergner, Bernhard Kuster, Götz Hensel, Jochen Kumlehn, Stefan Engelhardt, Ingo Heilmann, Ralph Hückelhoven

## Abstract

Rho-of-plant small GTPases (ROPs) are regulators of plant polar growth and of plant-pathogen interactions. The barley ROP, RACB, is involved in susceptibility towards infection by the barley powdery mildew fungus *Blumeria hordei* (*Bh*) but little is known about the cellular pathways that connect RACB-signaling to disease susceptibility. Here we identify novel RACB interaction partners of plant or fungal origin by untargeted co-immunoprecipitation of constitutively active (CA) RACB tagged by green-fluorescent protein from *Bh*-infected barley epidermal layers and subsequent analysis by liquid chromatography-coupled mass spectrometry. Three of immunoprecipitated proteins, a plant phosphoinositide phosphatase, a plant phosphoinositide phospholipase and a putative *Bh*-effector protein are involved in the barley-*Bh-*pathosystem and support disease resistance or susceptibility, respectively. RACB and its plant interactors bind to overlapping anionic phospholipid species *in vitro*, and in case of RACB, this lipid-interaction is mediated by its carboxy-terminal polybasic region (PBR). Fluorescent markers for anionic phospholipids show altered subcellular distribution in barley cells during *Bh-*attack and under expression of a RACB-binding fungal effector. Phosphatidylinositol 4-phosphate, phosphatidylinositol 3,5-bisphosphate and phosphatidylserine show a distinct enrichment at the haustorial neck region, suggesting a connection to subcellular targeting of RACB at this site. The interplay of ROPs with anionic phospholipids and phospholipid–metabolizing enzymes may thus enable the subcellular enrichment of components pivotal for success or failure of fungal penetration.

## Introduction

Although plant immunity is increasingly well understood, the understanding of mechanisms of disease susceptibility is still far from complete. Pathogen effectors target host proteins for suppression of plant immunity or trigger the function of plant negative regulators of immunity. Beyond this, some host susceptibility factors do not act in regulation of host immunity but serve the pathogenś demands in nutrition or in transforming host cell architecture (van Schie and Takken, 2014, Saur and Hückelhoven, 2021). Specific Rho-of-plant small monomeric GTPases (ROPs) might function as such susceptibility factors (Schultheiss *et al*., 2003). ROPs are the only representatives of the Rho-family of small GTPases that are found in plants (Yang, 2002) and, historically have also been termed RACs (rat sarcoma-related C botulinum substrate) (Winge *et al*., 2000). ROPs are considered as regulators of polar signaling processes in development, but can also regulate stress and immune responses (Kost *et al*., 1999, Jones *et al*., 2002, Akamatsu *et al*., 2013, Fratini *et al*., 2021). ROP signaling depends on the binding of ROPs to guanine nucleotides, and ROPs bound to GDP are signaling-inactive, whereas GTP-bound ROPs are signaling-competent (Berken, 2006). In some cases, the interaction between ROPs and downstream signaling proteins (here called executors) can be indirect and may require the presence of scaffolding proteins. Interactors of Constitutive Active ROPs (ICRs; also called ROP Interactive Partners (RIPs)) and ROP-Interactive and CRIB-Motif Containing Proteins (RICs) have previously been described as relevant ROP scaffolds (Wu *et al*., 2001, Lavy *et al*., 2007, Li *et al*., 2008, McCollum *et al*., 2020, Engelhardt *et al*., 2023). In *Arabidopsis thaliana* (*At*), RICs mediate ROP-effects on cytoskeletal organization, pavement cell morphogenesis or hormone responses (Wu *et al*., 2001, Gu *et al*., 2005, Fu *et al*., 2009, Choi *et al*., 2013, Lin *et al*., 2013, Zhou *et al*., 2015). Activated ROPs associate with the plasma membrane through C-terminal sequence motifs in the hypervariable region of the ROP proteins. In particular, a polybasic region (PBR) of positively charged lysine and arginine residues has been proposed to be essential for the interaction with membrane lipids *in vitro*, and for the recruitment of ROPs into anionic phospholipid-containing plasma membrane domains *in planta* (Platre *et al*., 2019; for fundamental work compare also Heo *et al*., 2006). For PM-interaction, type I ROPs are additionally cysteine-prenylated at the carboxyterminal CaaX-box motif and can be additionally and reversibly *S-*acylated (Sorek et al. 2017). Type II ROPs are constitutively *S*-acylated at their GC-CG-box (Yalovsky, 2015).

The membrane positioning of ROPs has been proposed to be governed in part by the presence of anionic phospholipids (Platre et al., 2019; Sternberg et al. 2021). Anionic phospholipid species constitute less than 1% of all lipids in plant leaves, but play vital roles in plant development and polar growth processes (Noack and Jaillais, 2020). Phosphoinositides (PtdIns), are an important group of anionic phospholipids that are formed by the phosphorylation of the membrane lipid phosphatidylinositol by specific lipid kinases at specific hydroxyl groups of the inositol head group. Reversely, PtdIns can be dephosphorylated by specific phosphatases or the inositol-phosphate headgroup can be cleaved from the diacylglycerol backbone by phospholipases (Mueller-Roeber and Pical, 2002, Zhong and Ye, 2003, Gerth *et al*., 2017). Another anionic phospholipid, phosphatidylserine (PtdSer), is generated by the addition of L-serine to phosphorylated diacylglycerol or via headgroup exchange from phosphatidylcholine or phosphatidylethanolamine with L-serine. With the help of genetically encoded cellular lipid markers and reverse genetics, a role for anionic phospholipids in stress responses, cellular trafficking processes and cell polarity was established (van Leeuwen *et al*., 2007, Gerth *et al*., 2017, Noack and Jaillais, 2020, Qin *et al*., 2020). Importantly, anionic phospholipids serve as targeting signals for proteins such as ROPs and their interactors (Platre *et al*., 2019, Kulich *et al*., 2020), and both PtdIns and PtdSer have been shown to influence the distribution of ROPs at plant membranes (Platre et al., 2019; Fratini et al., 2021). For instance, PtdSer-dependent recruitment of *At*ROP6 through its PBR is essential for ROP-dependent downstream processes during root gravitropism (Platre *et al*., 2019). Type II ROPs, too, require an intact PBR for lipid interaction and positioning in plasma membrane domains (Lavy and Yalovsky 2006; Sternberg et al. 2021).

A possible interplay of ROPs with anionic phospholipids during plant-pathogen interactions is less well established. The barley ROP, RACB, acts in signaling processes supporting susceptibility towards infection by the barley powdery mildew fungus *Blumeria hordei* (*Bh*) (Schultheiss *et al*., 2002, Schultheiss *et al*., 2003, Hoefle *et al*., 2011). Overexpression of constitutively activated RACB (RACB-CA) leads to more frequent fungal cell entry and formation of haustoria in intact plant cells that engulf the apoplastic fungal infection structure with an extrahaustorial matrix and an extrahaustorial membrane of host origin. Gene-silencing of *RACB* limits *Bh’*s penetration success and haustorial expansion, highlighting the role of RACB as a susceptibility factor (Schultheiss *et al*., 2002, Schultheiss *et al*., 2003, Hoefle *et al*., 2011). Interestingly, the signaling-active form of RACB appears to be required for its role as a susceptibility factor, as only overexpression of RACB-CA enhanced fungal infection success, whereas neither the overexpression of wild type (WT) RACB nor the dominant negative (DN) RACB increased barley susceptibility (Schultheiss *et al*., 2003). Several RACB-interacting executors and regulators are also involved in the barley-*Bh* pathosystem (Hoefle *et al*., 2011, Huesmann *et al*., 2012, Reiner *et al*., 2016, Trutzenberg *et al*., 2022) and three RACB-binding scaffold proteins, RIC157, RIC171 and RIPb, can support susceptibility towards *Bh*, when overexpressed. Furthermore, RIC157, RIC171 and RIPb interact with activated RACB at the PM *in planta*. During *Bh* attack, these ROP scaffolds localize to the haustorial neck region together with RACB (Schultheiss *et al*., 2008, McCollum *et al*., 2020, Engelhardt *et al*., 2023). PM localization of RACB appears to be essential for its role in susceptibility, since a truncated RACB mutant that lacks the prenylation motif “CSIL” mis-localizes to the cytosol and does not enhance fungal penetration (Schultheiss *et al*., 2003, Weiss *et al*., 2022). While the role of RACB as a susceptibility factor in the barley-*Bh* pathosystem is well supported, it remains unclear how RACB is subcellularly positioned and how the RACB signaling machinery may be recruited to the site of fungal entry at the PM.

Here, we identify three RACB interaction partners of plant and fungal origin, which establish a link between RACB and anionic phospholipids known to serve as position signals for protein-PM-recruitment. Anionic phospholipid markers display a distinct pattern of subcellular distribution during *Bh* attack, indicating a potential function in the barley-*Bh*-pathosystem. The identified RACB-interacting proteins as well as RACB all bind anionic phospholipids *in vitro*. RACB, binding to anionic phospholipids requires its intact PBR, and a RACB PBR substitution variant, RACB-5Q, defective in lipid binding *in vitro*, fails to localize to the PM *in planta* and is no longer signaling-competent with regard to barley susceptibility towards *Bh*. Together, the data indicate functional interplay between RACB and anionic phospholipids during barley/*Bh* interactions.

## Results

### PM localization is essential for RACB’s role as a susceptibility factor

To enable protein co-immunoprecipitation studies with tagged RACB, we generated stable transgenic barley lines overexpressing enhanced green fluorescent protein (eGFP)-tagged constitutively activated RACB, eGFP-RACB-CA, a RACB substitution variant lacking the four terminal amino acids “CSIL” of the CaaX-box prenylation site (eGFP-RACB-CA-ΔCSIL), or expressing free eGFP as control. We initially obtained 27 transformation events overexpressing eGFP-RACB-CA, 14 events for eGFP-RACB-CA-ΔCSIL and 17 for eGFP. After propagation of these independent events, offspring with the correct genotypes was identified by PCR, Western blotting and resistance against the selection marker hygromycin. Two independent transgenic lines for each construct were used for subsequent experiments. Using α-GFP Western blotting, we found that all fusion proteins were expressed in full size and could be enriched via α-GFP immunoprecipitation (IP; see Fig. 1a). The full-length eGFP-RACB-CA fusion localized to the PM of barley epidermal cells when analyzed by confocal laser scanning microscopy (CLSM). In contrast, the truncated eGFP-RACB-CA-ΔCSIL variant proved expectedly in the cytosol and the nucleus, similar to free eGFP (see Fig. 1b). Moreover, plants, overexpressing full-length eGFP-RACB-CA, were more susceptible towards haustorial invasion by *Bh*, when compared to plants overexpressing eGFP-RACB-CA-ΔCSIL or free eGFP (see Fig. 1c). This showed that the full length eGFP-RACB-CA fusion expressed in the transgenic barley lines was functional as a susceptibility factor. The observation that eGFP-RACB-CA-ΔCSIL did not mediate enhanced susceptibility is consistent with earlier findings that RACB-CA needs to be PM-localized to act (Schultheiß et al. 2003; Weiss et al. 2022).

**Figure 1:**
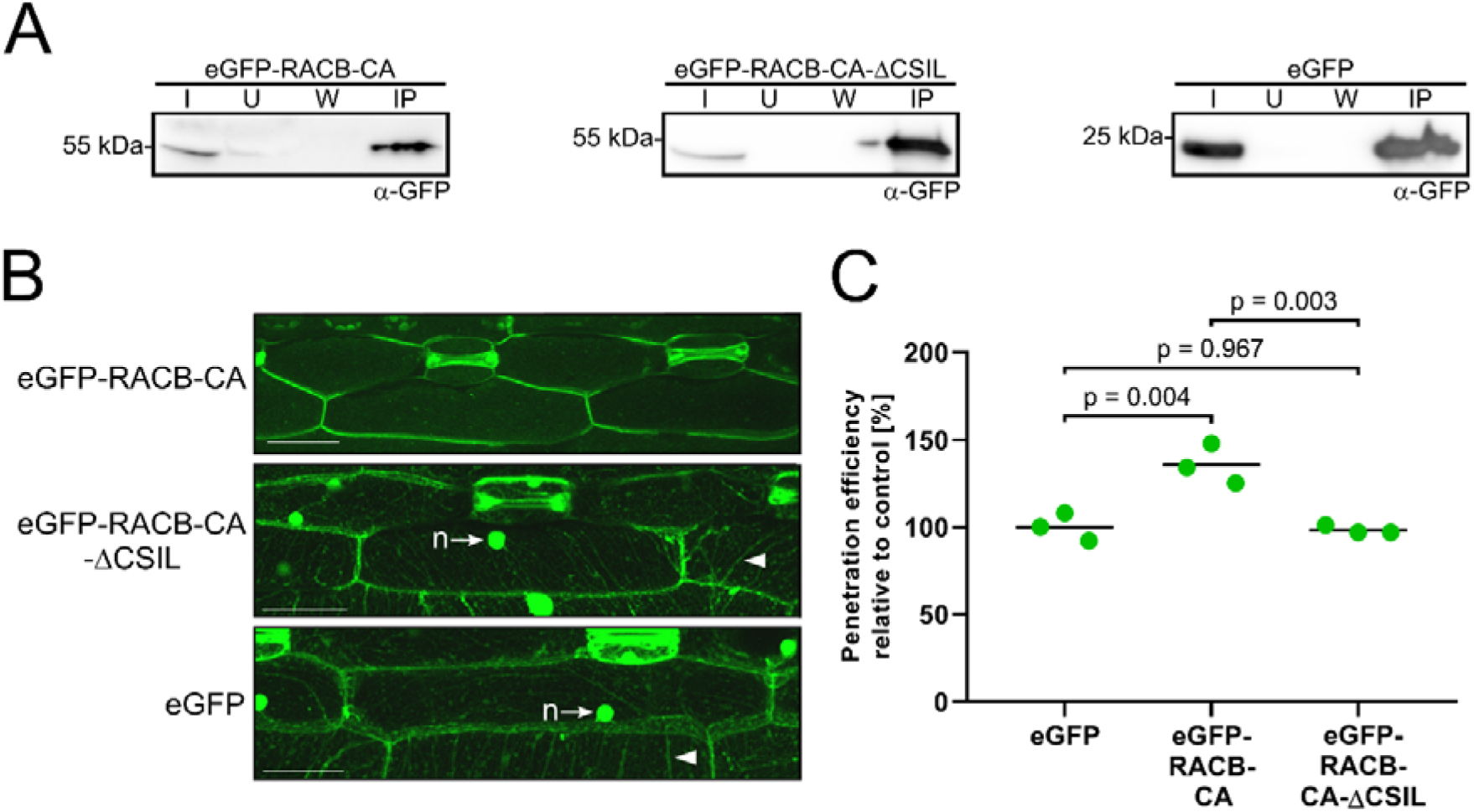
Transgenic barley plants overexpressing plasma membrane-localized eGFP-RACB-CA are super-susceptible towards *Bh* infection. **(a)**: eGFP and tagged fusion proteins show stable expression in first barley leaves and can be enriched via α-GFP IP. eGFP-tagged proteins were detected by α-GFP Western blotting. IP fractions: Input (I), Unbound (U), Wash (W) and enriched proteins (IP). **(b):** Full-length eGFP-RACB-CA localizes to the PM of barley epidermal cells, while eGFP-RACB-CA-ΔCSIL and free eGFP are visible in the cytosol (arrowhead: cytosolic strands) and nucleus (n, arrows). CLSM investigated subcellular localization. Scale bar: 50 µm. Images are maximum projections of at least 29 Z-steps of 2 µm increment optical sections. Image brightness was uniformly enhanced post-scanning for better visibility. **(c)**: Overexpression of full-length eGFP-RACB-CA renders barley plants more susceptible towards *Bh* invasion, whereas eGFP-RACB-CA-ΔCSIL and free eGFP show similar levels of lower susceptibility. For each construct, three primary leaves of one plant line were tested. Lines were: BG654 E2 for eGFP-RACB-CA, BG655 E1 for eGFP-RACB-CA-ΔCSIL and BG656 E1 for eGFP. Each data point represents the susceptibility of one leafthat was analyzed at 100 interaction sites. Crossbar depicts average susceptibility relative to the eGFP control. Statistical analysis was performed using a One-way ANOVA with Tukey’s honestly significant difference test.

### Co-immunoprecipitation identifies novel RACB-CA interactors

To identify novel RACB protein interaction partners, we performed an untargeted co-immunoprecipitation (CoIP) from transgenic eGFP-RACB-CA plants. eGFP-tagged proteins and associated putative interaction partners were co-immunoprecipitated at 24 h post inoculation (hpi) from *Bh*-infected (or mock-treated) epidermal peels from first leaves of transgenic barley, and proteins were identified by liquid chromatography-coupled tandem mass spectrometry (LC-MS/MS; see Fig. S1). We used epidermal peels for this screening, because *Bh* only colonizes the epidermal cell layer of barley. Overall, we detected 1399 unique proteins of plant and fungal origin in the LC-MS/MS analysis that were ranked according to statistically significant enrichment in eGFP-RACB-CA samples and the presence of unique peptides. Comparison of *Bh*-infected full-length eGFP-RACB-CA and free eGFP samples revealed 24 proteins that were significantly enriched with eGFP-RACB-CA (full dataset in Supplemental Table 1). Under mock-inoculation conditions, 17 proteins were significantly enriched in samples from eGFP-RACB-CA-expressing plants. In the infected samples, one protein of *Bh* origin stood out, as it was more than 60-fold enriched and found in all biological and technical replicates of each eGFP-RACB-CA and eGFP-RACB-CA-ΔCSIL and was annotated as a putative *Bh* effector protein (N1JEY6_BLUG1). We called this protein 9o9 for “found in 9 out of 9 replicates”. Among identified proteins of plant origin, we focused on two new possible interaction partners after comparing the curated dataset with literature data, because these proteins were annotated in the context of phospholipid metabolism. During mock conditions, a phosphoinositide phospholipase C-like protein was significantly enriched with eGFP-RACB-CA (Student’s t test p<0.01), whereas during infection a PtdIns phosphatase was exclusively enriched in samples from eGFP-RACB-CA-expressing plants (p<0.001). More detailed analysis of the corresponding proteins identified their nature as *Hordeum vulgare* phospholipase C1 (*Hv*PLC1, HORVU2Hr1G013730.2) and suppressor of actin (SAC) phosphoinositide phosphatase *Hv*SAC-like (HORVU4Hr1G077220.4) (supplemental figures Fig. S2-S3). The 9o9 protein shares a phylogenetic origin with classical *Bh* candidate secreted effector proteins (CSEPs) (Spanu et al., 2010; Pedersen et al., 2012), while it lacks a clearly predicted secretion signal.

### *In vivo* interaction between RACB and 9o9, PLC1 and SAC-like

To test whether the immunoprecipitated 9o9, PLC1 and SAC-like proteins are direct interaction partners of barley RACB, we investigated their ability to interact in targeted assays. For Förster-Resonance Energy Transfer Fluorescence Lifetime Imaging Microscopy (FRET-FLIM) in *Nicotiana benthaminana*, we expressed monomeric eGFP (meGFP) fused to the N-terminus of RACB-CA as a FRET donor, and co-expressed N-or C-terminal mCherry-fusions of 9o9, PLC1 and SAC-like as FRET acceptors. C-terminally mCherry-tagged glutathione *S*-transferase (Kulich *et al*.) and CRIB46, a ROP GTPase-interactive protein domain of the RACB-CA interaction partner RIC171 (Schultheiss *et al*., 2008, Trutzenberg *et al*., 2022), were used as negative and positive controls, respectively. In the FRET-FLIM analyses both mCherry-fusions of 9o9 decreased the GFP-lifetime of meGFP-RACB-CA, indicative of direct protein-protein interaction (Fig. 2a). C-terminally tagged PLC1-mCherry and N-terminally tagged mCherry-SAC-like also reduced the lifetime of meGFP-RACB-CA (Fig. 2b,c). The FRET-FLIM analyses indicate that 9o9 from *Bh*, and PLC1 and the SAC-like protein from barley can directly interact with RACB-CA *in planta*. While fusions of 9o9 to both C-terminal or N-terminal mCherry tags resulted in significantly reduced fluorescence lifetime of meGFP-RACB-CA, data for PLC1 or SAC-like were only significant for the C-terminal or N-terminal mCherry-fusion of PLC1 or SAC-like, respectively (Fig. 2b, c). These differences might result from protein-specific features of the fluorescence fusions and possibly indicate a weaker interaction of PLC1 or SAC-like with RACB-CA.

**Figure 2:**
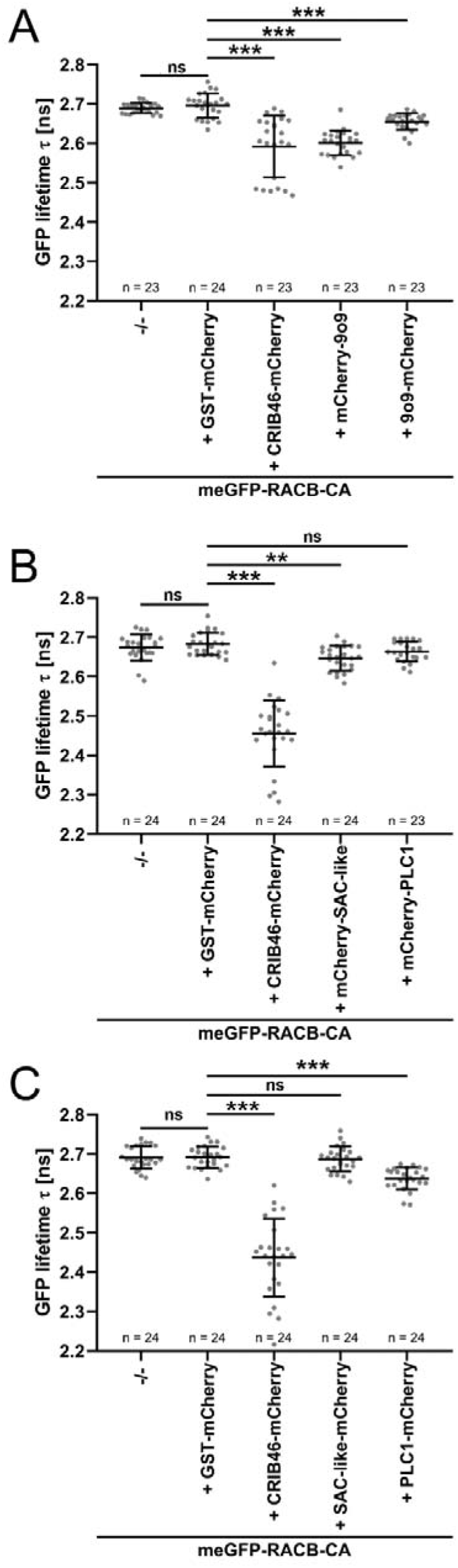
RACB-CA interacts with 9o9, PLC1 and SAC-like in FRET-FLIM experiments. (**a**) Co-expression of N-or C-terminally mCherry-tagged 9o9 decreased the fluorescence lifetime of GFP-RACB-CA, which indicates direct protein-protein interaction. (**b**) N-terminally tagged mCherry-SAC-like but not mCherry-PLC1 significantly reduced fluorescence lifetime of GFP-RACB-CA. (**c**) C-terminally tagged PLC1-mCherry but not SAC-like-mChrerry decreased lifetime of GFP-RACB-CA. GST-mCherry and CRIB46-mCherry were used as negative and positive controls, respectively (Schultheiss *et al*., 2008, Trutzenberg *et al*., 2022). FRET-FLIM measurements were conducted at the cell periphery of epidermal cells of transiently transformed *Nicotiana benthamiana* plants. Co-expression of fusion proteins was confirmed before measurements. All measurements were collected in three independent biological replicates. Number of observations (n) per construct is shown below each column. Crossbar and error bars show average and standard deviation. Statistical differences were calculated with Wilcoxon-Rank-Sum tests with Bonferroni-correction. ns p>0.05, **p <0.01, *** p< 0.001. -/-: FRET-donor-only control.

### 9o9 expression increases during *Bh* infection

To further characterize the three candidate RACB interactors in the context of *Bh* infection of barley plants, we investigated their expression by RT-qPCR (Fig. 3). A publicly available barley RNAseq dataset (Mascher *et al*., 2017, Rapazote-Flores *et al*., 2019) identified transcripts of *PLC1* and *SAC-like* in several barley tissues. Since both genes showed moderate expression levels in the epidermal cell layer that is colonized during fungal invasion, we analyzed whether *9o9*, *PLC1* or *SAC-like* were differentially expressed in epidermal cells upon *Bh* attack. Transcript abundance of *PLC1*, *SAC-like* and *9o9* was assessed in the barley epidermis at 6 h, 12 h and 24 h after *Bh* inoculation (Fig. 3). The *9o9* transcript increased over time, up to approx. 100-fold at 24 hpi compared to its initial transcript levels in spores and normalized to fungal tubulin gene expression (Fig. 3a). By contrast, the transcript abundance for the two plant genes did not strongly change upon *Bh* infection (Fig. 3b,c).

**Figure 3:**
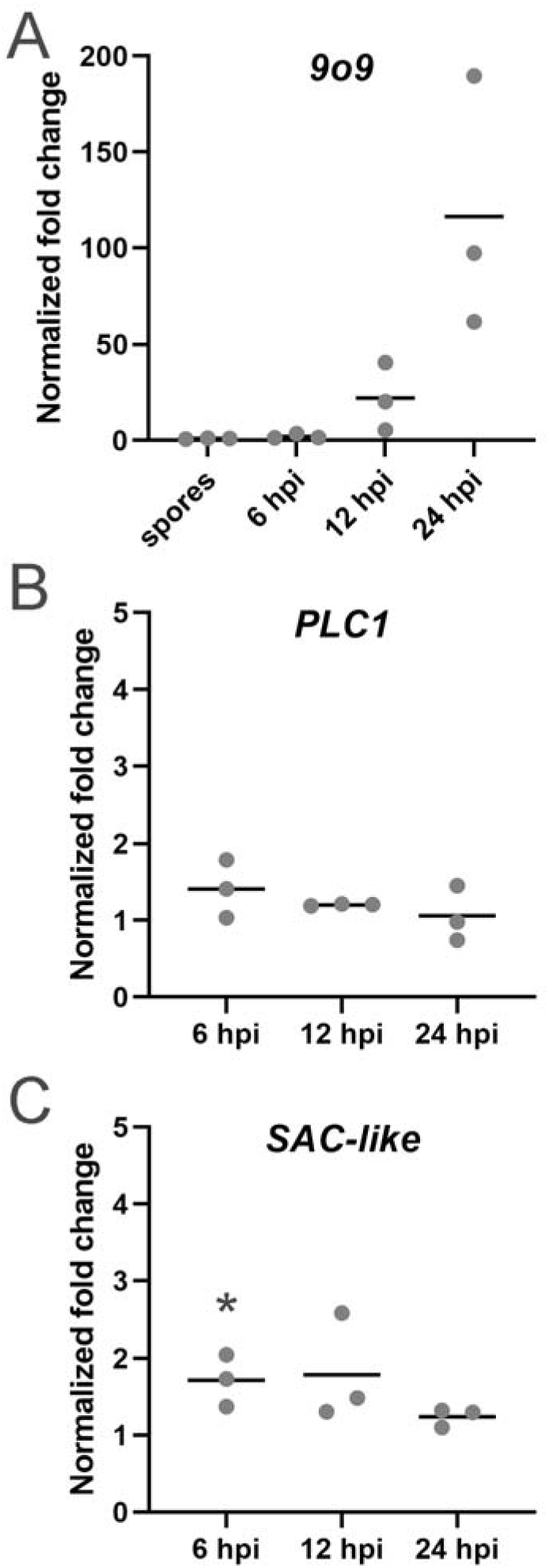
Gene expression of *Bh 9o9* increases during *Bh* invasion. The expression levels of *Bh 9o9* (**a**) and barley *PLC1* (**b**) and *SAC-like* (**c**) were measured during the early phase of *Bh* colonization at the indicated timepoints. Transcript of *9o9* was detectable in spores and increased during infection. Expression levels of *PLC1* and *SAC-like* only slightly differed during infection, but showed partially significant differences compared to their non-infected controls (as indicated by the p-values). Bars show reference gene-normalized fold changes in infected leaves, which were calculated according to using normalized gene expression in spores (**a**) or corresponding non-infected leaves (**b**, **c**) as a baseline. β*-TUB2* was chosen as a reference gene for *Bh*, while *UBC2* was taken as housekeeping genes for barley (Sherwood and Somerville, 1990, Schnepf *et al*., 2018). Bars show average fold-changes in normalized gene expression with standard deviation. Data was collected over three independent biological replicates. Statistical differences were calculated separately for each timepoint and gene, comparing the normalized expression of *PLC1* and *SAC-like* in infected leaves to that of corresponding non-infected leaves. Differences were assessed using *t-*tests with Holm-Sidak correction for multiple testing and only indicated in the graph when significant, *p<0.05.

### PLC1, SAC-like and 9o9 influence the barley-*Bh* interaction

The data indicated that the RACB-interactors, PLC1 and SAC-like, are expressed in barley tissues that are colonized by *Bh* and showed an upregulation of the *Bh* effector, 9o9, during the infection process. To test a possible role of the RACB interactors in the barley-*Bh* interaction, we transiently expressed these proteins in barley plants under the control of the cauliflower mosaic virus *35S* promoter, or we silenced expression using RNA interference (RNAi) before plants were inoculated with *Bh*. Overexpression of 9o9 in barley epidermal cells increased the relative barley susceptibility towards *Bh*-invasion on average by 36% over the susceptibility of empty vector controls, as determined by counting haustoria frequencies at 40 hpi in individual transformed cells that were attacked by *Bh*. By contrast, RNAi (host-induced gene silencing) against *9o9* did not influence the penetration efficiency of *Bh* (Fig. 4a, b). Conversely, RNAi against *PLC1* or *SAC-like* increased barley susceptibility towards *Bh* infection on average by 36% or 67%, respectively, compared to empty vector controls (Fig. 4c, d), whereas overexpression of PLC1 or SAC-like did not change the susceptibility of barley against *Bh*. Data indicate that 9o9, PLC1 and SAC-like have opposing roles in the barley*-Bh* interaction, with ectopic fungal 9o9 expression promoting susceptibility and barley PLC1 and SAC-like limiting fungal penetration success.

**Figure 4:**
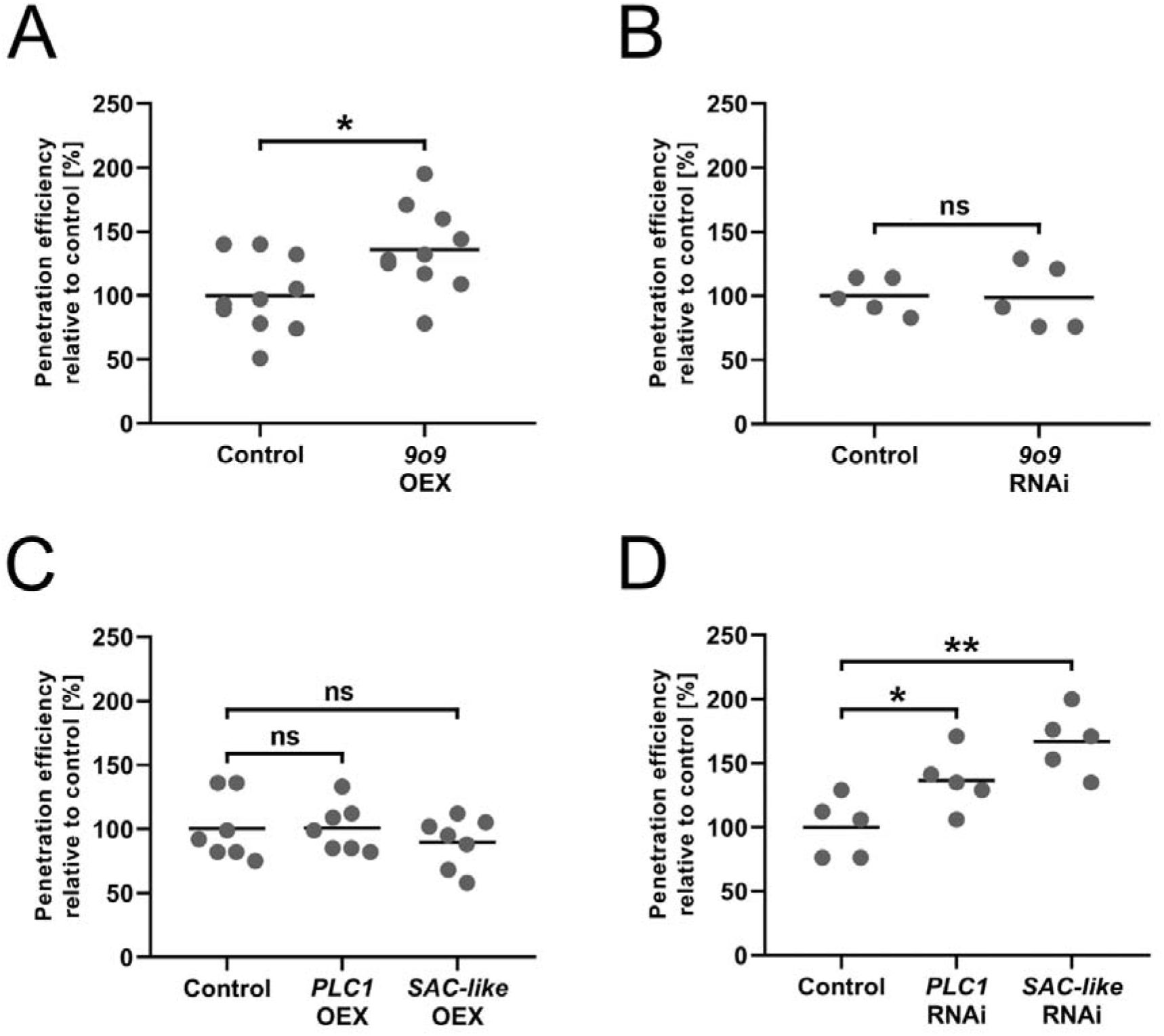
Single cell overexpression or silencing of *9o9, PLC1* and *SAC*-like influence in the barley-*Bh* pathosystem. The susceptibility of barley epidermal cells towards penetration by *Bh* was investigated after transient overexpression (**a**, **c**) or RNAi-mediated silencing (**b**, **d**) of 9o9, PLC1 and SAC-like at 40 hpi. The respective empty vectors were used as controls. Overexpression of 9o9 and silencing of *PLC1* and *SAC-like* lead to increased susceptibility, whereas silencing of *9o9* and overexpression of PLC1 and SAC-like showed no influence. RNAi-silencing specificity was confirmed using the si-Fi RNAi-off-target prediction tool (Lück *et al*., 2019). Each datapoint shows the *Bh* penetration efficiency of a single experiment relative to its averaged empty vector control. Crossbars display the average susceptibility from ten (A), five (B, D) and seven (C) independent biological experiments. Statistical differences were calculated with two-tailed Student’s *t*-tests comparing an overexpression or silencing construct with its respective empty vector control: ns p>0.05, * p<0.05, **p <0.01. OEX: overexpression.

### RACB, PLC1 and SAC-like show overlapping patterns of lipid-binding capabilities

The requirement of PM-association of RACB (cf. Fig. 1) as well as the interaction of RACB with PLC1 or SAC-like proteins (cf. Figs. 2) suggested a link between RACB function and anionic phospholipids. Like other ROPs (Platre et al., 2019), RACB contains a PBR predicted to bind anionic phospholipids. Computational analysis suggested that also PLC1 and SAC-like can interact with phospholipids and potentially possesses enzymatic activity against phosphoinositides. To investigate which phospholipids might bind to the 9o9, PLC1 or SCA-like proteins, *in vitro* lipid-binding assays were performed with recombinantly expressed GST-or MBP-tagged proteins (Fig. 5). A GST-tagged RACB-WT protein was included in these experiments that is similar to *Arabidopsis thaliana At*ROP6 that has previously been shown to display lipid-binding capacity (Platre *et al*., 2019). In protein-lipid-overlay experiments, GST-RACB-WT and its MBP-tagged PLC1 and SAC-like interactors displayed similar lipid-binding features and bound to PtdIns-monophosphates (PtdIns3P, PtdIns4P and PtdIns5P) (Fig. 5). In a GST-RACB-5Q substitution variant (RACB-K184Q, K185Q, K186Q, K187Q, K188Q), the substitution of five lysine residues in the PBR for uncharged glutamine (Q) residues fully abolished lipid binding (Fig. 5).

**Figure 5:**
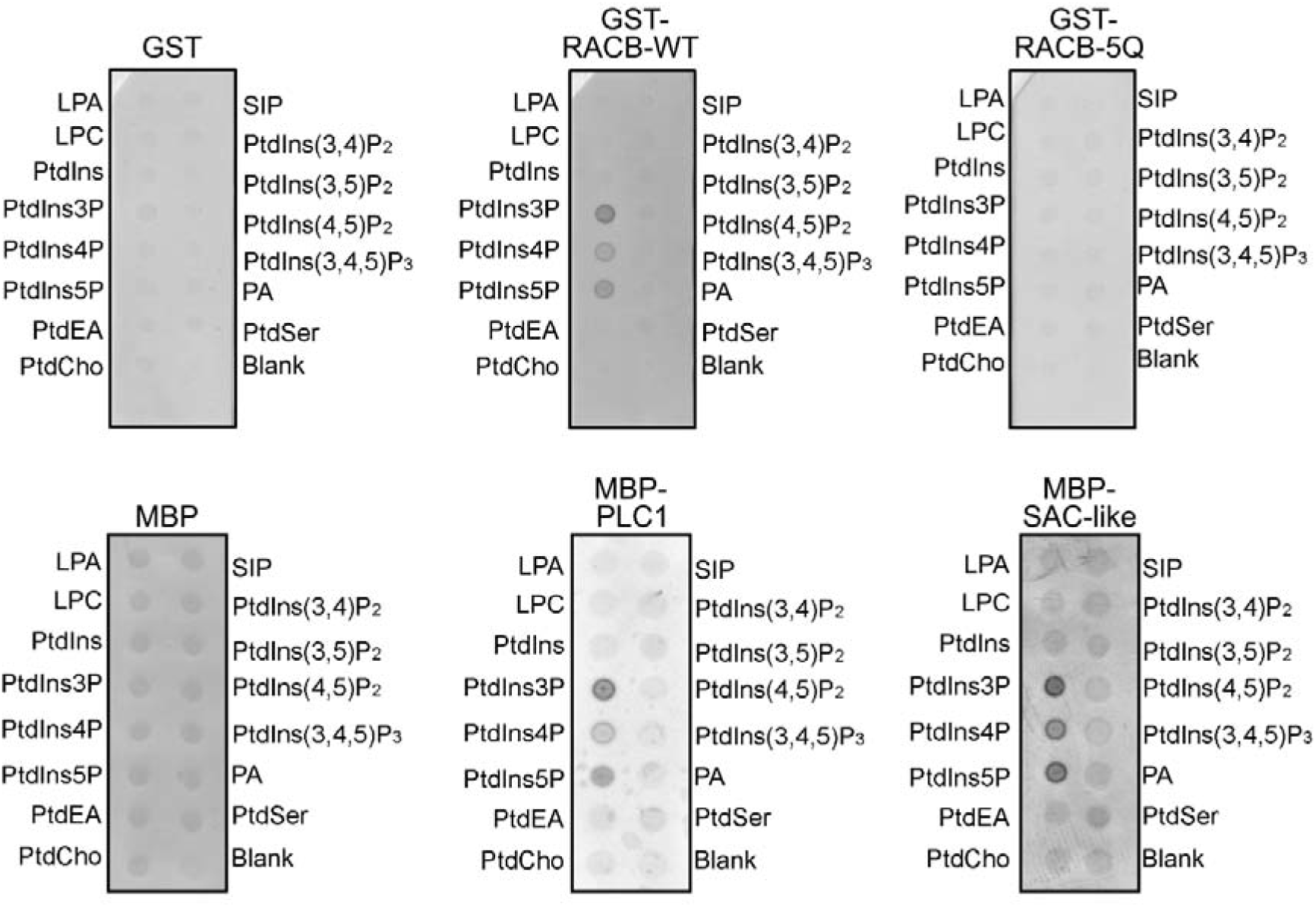
RACB and its candidate interactors associate with PtdIns-monophosphates *in vitro*. RACB and its candidate interactors show lipid-binding in *in vitro* protein-lipid-overlay experiments. Recombinant proteins were purified from *E. coli* using affinity chromatography. Recombinant proteins were incubated with lipid-spotted membranes and detected via α-GST or α-MBP antibodies. Colored dots indicate association with the respective lipid species. Free GST and MBP were used as non-lipid-binding controls. LPA: lysophosphatidic acid; LPC: lysophosphatidylcholine; SIP: sphingosine 1-phosphate; blank: no lipid spotted.

### RACB’s PBR is essential for its signaling function and membrane localization

Since the RACB-5Q substitution variant did not bind phospholipids, we investigated whether PBR mutants were functional *in planta*. As the PBR of the related AtROP6 has previously been shown to mediate membrane-association of AtROP6 (Platre *et al*., 2019), we first compared the subcellular localization of a GFP-RACB-5Q fusion with that of GFP-RACB-CA in barley epidermal cells. We further investigated whether the RACB-CA-5Q variant still interacted with its downstream partner, the canonical RACB-interactor and ROP-scaffold, RIC171. Using CLSM of transiently transformed barley epidermal cells (Fig. 6), soluble CFP, GFP and mCherry-RIC171 fusions, expressed as controls, displayed cytoplasmic and nucleoplasmic fluorescence distributions (Fig. 6a, top panels). By contrast when GFP-RACB-CA was co-expressed instead of the GFP control, both GFP-RACB-CA and mCherry-RIC171 signals were detected at the PM and for mCherry-RIC171 additionally in the nucleus (Fig. 6a, mid panels). The data are consistent with the previous report that PM associated RACB-CA can recruit RIC171 to the cell periphery (Schultheiss et al. 2008; Trutzenberg et al. 2022). By contrast, GFP-RACB-CA-5Q mislocalized to the cytoplasm and nucleoplasm (Fig. 6a, bottom panel). Moreover, in cells co-expressing GFP-RACB-CA-5Q and mCherry-RIC171, mCherry-RIC171 was not detected at the cell periphery but remained in the cytoplasm and nucleoplasm (Fig. 6a, bottom panels) in a similar distribution as was observed upon coexpression of mCherry-Ric171 with the controls, GFP and mCherry (Fig. 6a, top panels). As in yeast-2-hybrid experiments (Fig. 6b) and in FRET-FLIM experiments (Fig. 6c), RACB-CA-5Q interacted with RIC171 in a similar manner as the parental RACB-CA, we conclude that lipid binding through the PBR is required for PM-association of RACB-CA and for its capability to co-recruit the ROP scaffold RIC171 to the cell periphery upon infection.

**Figure 6:**
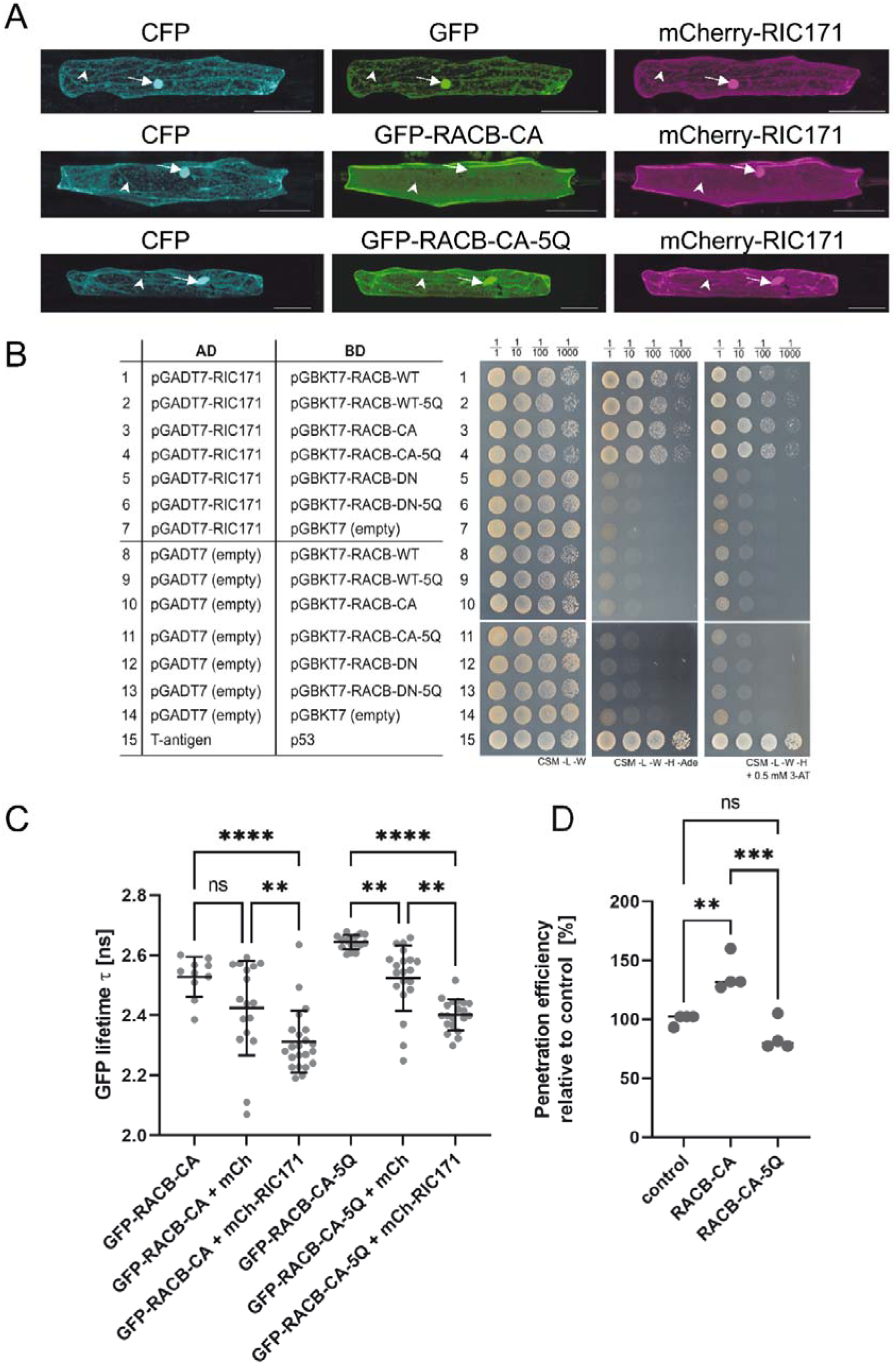
RACB-CA localization and function depend on positively charged amino acid residues in its PBR. **(a)** The subcellular localization of GFP-tagged RACB-CA or its derivate RACB-CA-5Q together with its mCherry-tagged putative downstream scaffold protein RIC171were investigated via CLSM after transient transformation of epidermal cells. Size bars represent 50µm. **(b)** Yeast two hybrid assay of Binding Domain (Vossen *et al*.) tagged RACB-variants (WT, wild type; CA, constitutively activated; DN, dominant negative) with Activation Domain (AD) tagged RIC171 in yeast on protein interaction non-selective complete supplement medium (CSM) plates lacking leucine and tryptophan (-L -W) or protein interaction selective plates lacking leucine, tryptophan, histidine and adenine (-L -W -H -Ade), or protein interaction selective plates supplemented with 0.5 mM 3-aminotriazole (-L -W -H + 0.5 mM 3-AT). Interaction indicative growth is visible for combinations between RIC171 and each of RACB-WT, RACB-WT-5Q, RACB-CA and RACB-CA-5Q. Interaction between T-antigen and p53 served as a positive control for the yeast two-hybrid experiment. **(c)** Interaction of GFP-tagged versions of RACB-CA with mCherry-RIC171 in FRET-FLIM experiments. Barley epidermal cells were transiently transformed via particle bombardment with expression constructs for either GFP-RACB-CA or GFP-RACB-CA-5Q as FRET-donors and mCherry-RIC171 as a FRET-acceptor. Free mCherry served as no-interaction controls. FRET-FLIM measurement were conducted at the cell periphery of the equatorial plane of barley epidermal cells 2 d after transformation. Stars display statistically significant differences between the samples (Tukey test, ns p>0.05, **p <0.01, *** p< 0.001 based on an ANOVA (significant at α= 0.05). Measured cells were collected in at least 3 independent biological replicates. **(d)** The susceptibility of barley epidermal cells towards penetration by *Bh* was investigated after transient overexpression of RACB-CA or RACB-CA-5Q or an empty vector control, respectively. Each data point shows the *Bh* penetration efficiency of a single experiment relative to its averaged control. Crossbars display the average susceptibility from four independent biological experiments. Statistical differences were calculated with Tukey test, ns p>0.05, **p <0.01, *** p< 0.001 based on an ANOVA (significant at α= 0.05). control= empty vector transformation.

Finally, we tested if the RACB-CA-5Q substitution variant is still functional as a susceptibility factor in the barley-*Bh* pathosystem. Using transient overexpression of regular RACB-CA or RACB-CA-5Q in barley epidermal cells, we detected that the expression of RACB-CA significantly enhanced relative fungal penetration success by 38% compared to empty vector controls (Fig. 6d). By contrast, the expression of RACB-CA-5Q reduced fungal penetration success by 15% of control levels. While this decrease was statistically not different from penetration success in empty vector controls, it was significantly lower than that observed upon expression of RACB-CA (Fig. 6d). In summary, these experiments showed that the RACB-CA-5Q mutant can still interact with downstream partners, such as RIC171. However, both the subcellular localization and the function of RACB-CA in disease susceptibility are compromised by the substitutions in the PBR that abolish PtdIns-binding.

### Some anionic phospholipids localize to the site of haustorial invasion by *Bh*

Since RACB bound to some anionic phospholipids *in vitro* and this lipid binding capability was found to be relevant *in vivo*, we next used fluorescent lipid biosensors to monitor the subcellular distribution of anionic phospholipids, first in untreated barley epidermal cells and then during *Bh*-infection. We chose biosensors for PtdIns4P, PtdIns(3,5)P_2_, PtdIns(4,5)P_2_ or PtdSer for subcellular localization studies, because these lipids had either been linked to ROP-signaling before or were shown to display an altered behavior during pathogen attack in other plants (Kost *et al*., 1999, Hirano *et al*., 2018, Platre *et al*., 2019, Qin *et al*., 2020, Fratini *et al*., 2021). Importantly, in previous reports PtdIns(4,5)P_2_ has been described as a susceptibility factor for other plant-pathogen interactions (Shimada *et al*., 2019, Qin *et al*., 2020). For *in planta* visualization of the different phospholipids we used fluorophore-tagged protein domains that bind specific phospholipids (Simon *et al*., 2014, Hirano *et al*., 2017). These biomarkers were used for transient transformation of barley epidermal cells that were then observed via CLSM either in non-infected controls (Fig. 7a) or during fungal invasion at 16-20 hpi (Fig. 7b-e). All biomarkers had to be imaged with individual laser excitation levels and detector gain to detect fluorophore-specific signals. A GFP-2xPH^FAPP1^ biosensor for PtdIns4P, bound by a double pleckstrin homology (PH) domain of the human Four-Phosphate Adapter Protein (2xPH^FAPP1^, Simon *et al*. (2014)), decorated the PM of non-infected barley epidermal cells when compared to mCherry as a cytoplasmic marker (Fig. 7a). The GFP-2xPH^FAPP1^ marker also labeled the PM also upon *Bh*-attack (Fig. 7b). Fluorescence patterns were detectable even though autofluorescence was usually strong in cell wall appositions at sites of attack and overlapping with fluorophore signals (compare Fig 7b and 7f for a pure autofluorescence signal from an attacked but non-transformed neighbor cell). However, we additionally detected a clear enrichment of the GFP-2xPH^FAPP1^ marker at the haustorial neck-region of successfully *Bh-*colonized cells, with little autofluorescence at such sites (Fig. 7b, see supplemental figure S4 for higher magnification). At sites of successful penetration, signals were strong and originated from the emission λ-spectrum of the GFP-2xPH^FAPP1^ -derived GFP signal. Interestingly, the PtdIns4P marker was not detected at the extrahaustorial membrane that is in continuum with the plant PM but of a distinct composition (Kwaaitaal *et al*., 2017).

**Figure 7:**
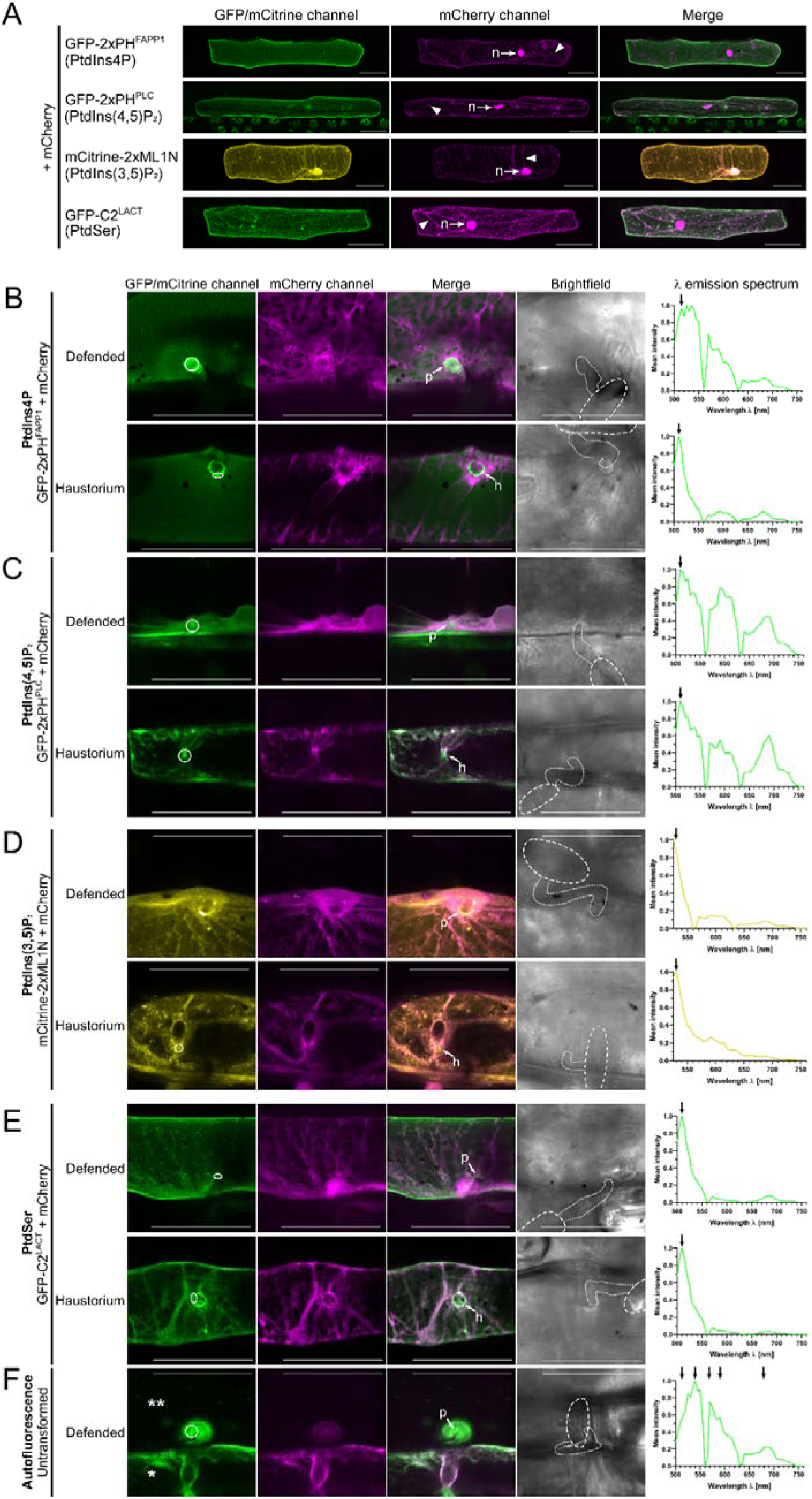
Subcellular localization of PtdIns4P, PtdIns(4,5)P_2_, PtdIns(3,5)P_2_, and PtdSer in unstressed and Bh-infected barley epidermal cells. **(a)** The subcellular localization of four anionic phospholipid species was investigated in barley via CLSM after transient transformation of epidermal cells with lipid-species-specific genetically encoded biomarkers (Simon *et al*., 2014, Hirano *et al*., 2017, Platre *et al*., 2018). All four species showed at least partial localization to the PM. Free mCherry was co-transformed as a marker highlighting localization in the cytosol (arrowhead: cytosolic strands) and nucleus (n, arrows). All images show Z-stack maximum intensity projections composed of at least 16 XY-optical sections captured in 1.5 µm Z-steps. Scale bar: 50 µm. Representative images from at least two independent biological replicates are shown. Image brightness was uniformly enhanced post-scanning for better visibility. **(b-f)** Images of infected cells were taken between 16-20 hpi. High magnification images of non-penetrated (“Defended”) and *Bh-*colonized (“Haustorium”) cells are shown. Arrows point to unsuccessful penetration attempts (p) or haustorial entry points (h). PtdIns4P **(b)** and PtdSer (**e**) were enriched at the haustorial neck region of *Bh-*colonized cells. Signal from PtdIns(4,5)P_2_ (**c**) was primarily visible in cytosol and nucleus, but was hard to evaluate due to very low fluorescence levels. PtdIns(3,5)P_2_ **(d)** was slightly enriched at the cell wall apposition of non-penetrated cells. The circles in the GFP/mCitrine-images show regions-of-interest that were λ-scanned to evaluate fluorescence emission spectra. λ-scanning was performed in 5 nm detection steps after excitation with a 488 nm (GFP) or 514 nm (mCitrine) laser source. Only in non-penetrated GFP-2xPH^FAPP1^-transformed cells and all GFP-2xPH^PLC^-transformed cells a strong presence of non-GFP-signals could be detected that was likely auto-fluorescence emitted by phenolic compounds released during plant defense responses. In all other cells spectra matched that of GFP or YFP (fluorophore emission maxima are highlighted by arrows;). For reference, **(f)** shows both autofluorescence in an untransformed, *Bh-*attacked cell (**) and, side by side, a single transformed cell (*) imaged with the settings for PtdIns(4,5)P_2_. λ-scanning was performed in the indicated region of interest (circle) to characterize auto-fluorescence emitted during high laser excitation levels and detector gain. Note the lack of the typical GFP peak around 510 nm and the peaks/shoulders from autofluorescence at about 540, 570, 590 and 690 nm. Free mCherry was co-transformed as a marker for cytosolic and nuclear fluorescence. All images except brightfield pictures show Z-stack maximum intensity projections of at least 4 XY-optical sections captured in 1.5 µm Z-steps. Brightfield images show single XY-optical sections, in which spores are outlined with dashed lines and appressoria are highlighted with dotted lines. Scale bar: 50 µm. Representative images from at least five events in each at least two independent biological replicates are shown. All images were taken with individual laser excitation strength and detector gain for higher image clarity. Image brightness was uniformly enhanced post-scanning for better visibility.

A biosensor for PtdIns(4,5)P_2_, a GFP-fused double PH-domain of rat PLCδ1 (2xPH^PLC^, Simon *et al*. (2014)), showed a prominent localization at the PM in non-infected cells, with some background signals coming from the cytosol and nucleus (Fig. 7a). By contrast, in *Bh*-attacked cells, the PM-associated intensity of the GFP-2xPH^PLC^ marker was weaker and became hard to determine, especially due to the intrinsic autofluorescence (Fig. 7c). The reduced PM-fluorescence was accompanied by an increased cytoplasmic signal for GFP-2xPH^PLC^ (Fig. 7c), consistent with a reduction in PtdIns(4,5)P_2_ at the PM upon *Bh*-attack.

The distribution of PtdIns(3,5)P_2_ was tracked by mCitrine fused to a tandem repeat of the lipid-binding domain of mammalian Mucolipin 1 (Hirano *et al*., 2017). The mCitrine-2x-MLN1 marker was visible at the cell periphery, in the nucleus and in the cytoplasm of non-infected cells (see Fig. 7a). While it is difficult to discern from our imaging setup, mCitrine-2x-MLN1 distributed in a pattern differing somewhat from PM localization. Moreover, in some cells mCitrine-2x-MLN1 fluorescence was also detected in endosomal vesicle-like structures (Fig. 7d). As PtdIns(3,5)P_2_ has been reported to reside at the plasma membrane, endosomes and tonoplast (Hirano el al. 2017; Hirano el al. 2018; Bak *et al*., 2013, Nováková *et al*., 2014), the observed pattern might represent those localization. During *Bh* interaction, the mCitrine-2x-MLN1-marker was present but little distinct at sites of haustorium accommodation and seemingly a bit more enriched around non-penetrated plant cell wall appositions (Fig. 7d).

The PS-marker, a GFP-tagged C2-domain of bovine Lactadherin (C2^LACT^, Platre *et al*. (2018)), was detected mainly at the PM, but also weakly in the cytosol of non-infected barley epidermal cells (Fig. 7a). In most cases, the PS-marker was also visible in tiny, potentially vesicular speckles. During *Bh-*infection, GFP-C2^LACT^ remained at the PM, but still exhibited a stronger cytoplasmic background signal (Fig. 7e) in a similar pattern as the GFP-2xPH^PLC^ marker for PtdIns(4,5)P_2_. In *Bh-*colonized cells, a localization at the haustorial neck-region was evident for the PS-marker that was more distinct at this position than the cytosolic mCherry signal (Fig. 7e, see supplemental figure S4 for higher magnification). Similar to the GFP-2xPH^FAPP1^ marker for PtdIns4P, the GFP-C2^LACT^ marker was also not enriched at the extrahaustorial membrane. In conclusion, the distribution of the various fluorescent biosensors indicate a potential involvement of anionic phospholipids in the barley-*Bh* interaction, as three of the four lipid biosensors tested (for PtdIns4P, PtdIns(4,5)P_2_ and for PtdSer) showed an altered localization or subcellular distribution during fungal attack.

Because 9o9 was originally found in nine Co-IPs with RACB-CA and lipid-modifying PLC1 and SAC-like, we wondered whether 9o9 could change the subcellular pattern of the tested lipid markers. We found an effect of 9o9 on the PtdIns(3,5)P_2_ marker mCitrine-2x-MLN1. Co-expression of 9o9 apparently not only supported fungal penetration but also supported the presence of mCitrine-2x-MLN1 at the haustorial neck region (Fig. 8).

**Figure 8:**
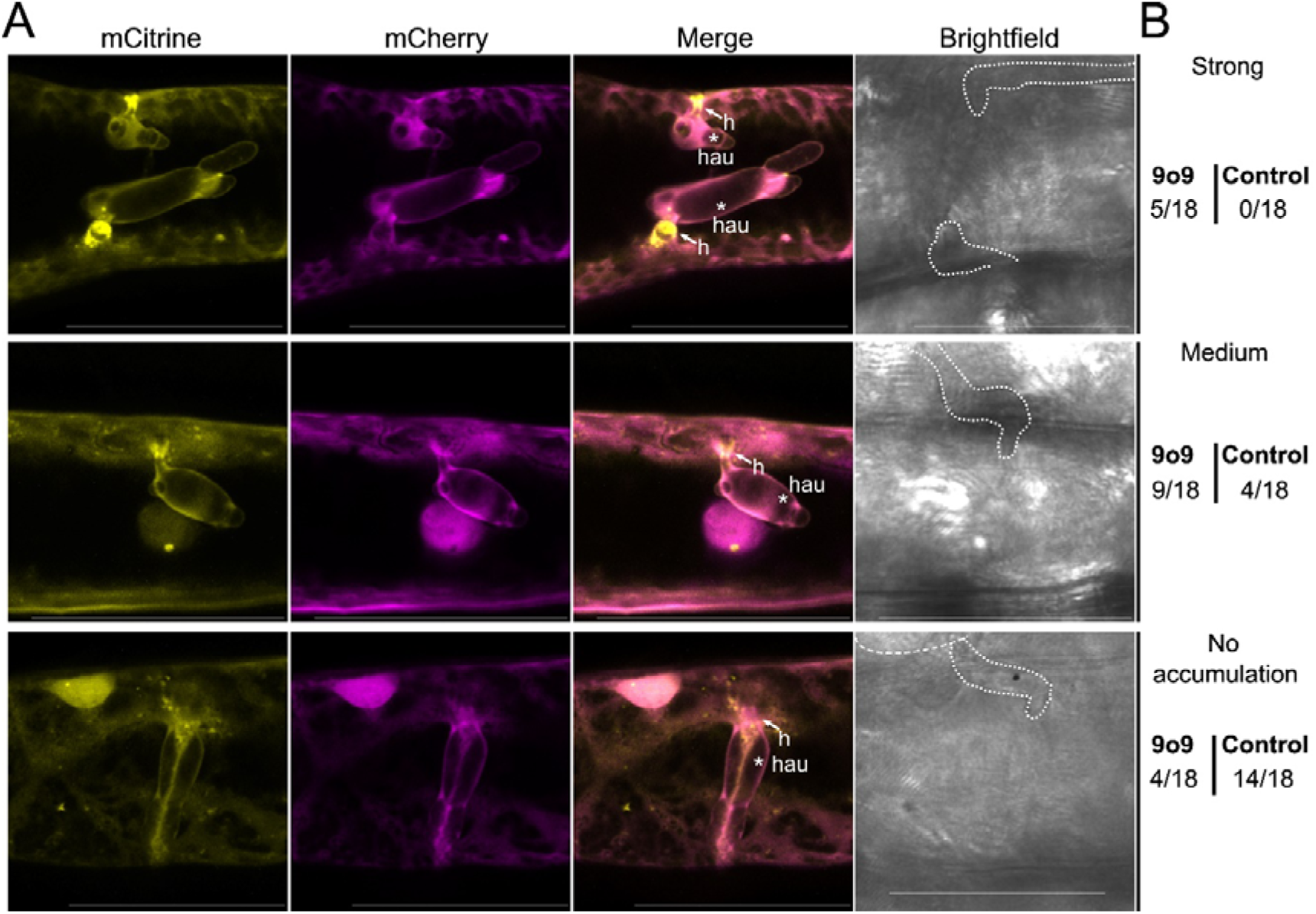
Effect of 9o9 on the subcellular localization of PtdIns(3,5)P_₂_ marker in *Bh*-infected barley epidermal cells. (a) The subcellular localization of the anionic phospholipid PtdIns(3,5)P_₂_ was analyzed in barley epidermal cells via confocal laser scanning microscopy following transient co-transformation with *Bh* 9o9 and the PtdIns(3,5)P_₂_-specific biosensor Citrine-2x-ML1N (Hirano et al., 2017). Free mCherry was co-transformed as a cytosolic marker. Images of infected cells were taken between 16–20 hpi. High-magnification images of *Bh*-colonized cells are shown; haustorial entry points are indicated with (h) and the asterisk (*) indicates the haustorial body. All images show Z-stack maximum intensity projections composed of at least 16 XY-optical sections captured in 1.5 μm Z-steps. Brightfield images represent single sections of the transmission channel, with the haustorium out of focus. Scale bar: 50 μm. Image brightness was uniformly enhanced post-acquisition for better visualization. Representative images illustrate three categories of PtdIns(3,5)P_₂_ accumulation at haustorial entry sides/neck regoin: strong accumulation (upper panel), medium accumulation (middle panel), and no accumulation (lower panel). (b) Counting of PtdIns(3,5)P₂ accumulation patterns observed in each eighteen cells expressing 9o9 or control cells co-transformed with the empty vector. In 9o9-expressing cells, PtdIns(3,5)P_₂_ accumulation at haustorial entry points was seen in the majority of cells. In most controls cells, PtdIns(3,5)P_₂_ accumulation was absent or weak.

To gain first hints towards a possible mechanism of 9o9 action, we wanted to test whether 9o9 indeed preferentially interacts with activated RACB-CA when compared to wild type RACB-WT or dominant negative RACB-DN. In targeted co-immunoprecipitation experiments, we co-expressed HA-tagged HA-9o9 or 9o9-HA with GFP-RACB-CA, GFP-RACB-WT or GFP-RACB-DN in *Nicotiana benthamiana.* GFP-RACB-DN was not stably expressed in *N. benthamiana* but free GFP, GFP-RACB-CA, GFP-RACB-WT were well detectable in Western blots after precipitation with GFP antibodies. HA-tagged versions of 9o9 co-precipitated with GFP-RACB-CA but neither with GFP-RACB-WT nor free GFP. This supports that 9o9 preferentially interacts with GTP-loaded RACB (supplemental Fig. S5). Protein complex modelling of protein dimers of GTP-loaded RACB with PLC1, SAC-like and 9o9 further suggests that all three interaction partners of RACB share an overlapping interface of protein-protein interaction with partially identical amino acids around the ROP effector loop of RACB responsible for binding the respective partner (supplemental Fig. S5).

## Discussion

### PM-association through interaction with PtdIns-monophosphates is essential for RACB function in disease susceptibility

ROPs need to associate with the PM for activation and downstream signaling activities in plant development (Yalovsky, 2015, Platre *et al*., 2019). Here we analyzed, how the barley RACB protein interacts with the PM, whether it binds membrane phospholipids, and whether this interaction is relevant for its role as a susceptibility factor in the barley-*Bh*-interaction. First, we confirmed previous findings with untagged or HA-tagged versions of RACB that overexpression of full-length PM-localized GFP-RACB-CA increased susceptibility towards *Bh* infection, whereas a mis-localized cytoplasmic GFP-RACB-CA-ΔCSIL mutant did not support susceptibility (Fig. 1b, C; (Schultheiss *et al*., 2003, Weiss *et al*., 2022)). Mechanistically, the RACB-CA-ΔCSIL mutant could be affected in its ability to associate with downstream executors, since these interactions are often observed at the PM (Schultheiss *et al*., 2008, Hoefle *et al*., 2011, McCollum *et al*., 2020). Alternatively, it is possible that cytoplasmic RACB-CA-ΔCSIL can interact with downstream executors but this interaction is spatially separated from usual ROP signaling processes at the PM. Irrespectively, both the subcellular localization of RACB and its activation status appear essential for RACB function in susceptibility. The ability to interact with phospholipids can also influence the subcellular localization and associated signaling capacities of ROPs, such as the type I ROP *At*ROP6 or the type II ROP *At*ROP11 (Platre *et al*., 2019; Sternber *et al*. 2021). *At*ROP6, a type I ROP similar to RACB, associates with PtdSer and with PtdIns-monophosphates (Platre *et al*., 2019). *In planta* PtdSer is essential for enrichment of *At*ROP6-CA in PM-nanodomains (Platre *et al*., 2019). By contrast, *At*ROP6-7Q did not cluster in nanodomains, accompanied by abolished signaling capacity (Platre *et al*., 2019). Data presented here indicate that barley RACB also interacts with different PtdIns-monophosphates *in vitro* (Fig. 5), and this lipid interaction was important for subcellular localization at the PM (Fig. 6a) and for the role of RACB as a susceptibility factor during the barley-*Bh*-interaction (Fig. 6d). Our experiments show that an interaction between ROPs and PtdIns-monophosphates can support compatible plant-pathogen interactions. Additionally, it was shown earlier, that the PBR of ROPs is also required for efficient prenylation of CaaX box motifs (Caldelari et al. 2001), such that failure of RACB-CA-5Q may arise from a combination of effects. In analogy to *At*ROP6 (Platre et al., 2019), activated barley RACB might enrich in PM-domains in a phospholipid-dependent manner. As the lipid binding-deficient RACB-CA-5Q protein could still interact with the scaffold protein RIC171 in yeast and *in planta*, most likely protein mis-localization rather than protein mis-folding was responsible for the failure of RACB-CA-5Q to recruit signaling partner proteins to the PM, or to function as a susceptibility factor. Our data indicate that the function of RACB as a susceptibility factor requires PM-localization, which is mediated by both CSIL prenylation (Schultheiss et al. 2003, Weiss et al. 2022), and the binding to PtdIns-monophosphates and possibly other anionic phospholipids through positively charged lysine residues in its PBR.

### 9o9 is a *Bh* effector protein targeting activated RACB

The untargeted RACB-CoIP from *Bh*-infected epidermal peels of barley plants identified 9o9, a putative effector protein from *Bh*, as a candidate RACB-CA interaction partner, and the 9o9-RACB-CA interaction could be verified by a range of other methods. The *Bh* 9o9 protein belongs to the CSEP superfamily of *Blumeria* spec. effectors (Fig. S3a; Pedersen et al. 2012) that contains many validated effector proteins with functions in effector-triggered susceptibility and effector-triggered immunity. However, with 317 amino acids in length, 9o9 is more prominent in size when compared to most characterized *Blumeria* effector proteins. *In planta* interaction of 9o9 with RACB-CA (Fig. 2a; Fig. S5) might be part of a potential virulence mechanism for 9o9, since exploiting susceptibility factors is a virulence strategy for several pathogens (Mackey *et al*., 2002, He *et al*., 2018, Choi *et al*., 2021). The expression of the *9o9* gene is strongly upregulated during the early stages of *Bh*-invasion (Fig. 3a), and transient overexpression of 9o9 in barley increased susceptibility towards *Bh* invasion (Fig. 4a). Our effort to prove physiological function of 9o9 by host-induced gene silencing were not successful. This may be explained by redundancy of 9o9 and other similar effectors or dissimilar effectors potential targeting of the same host pathway. This seems possible because *B. hordei* actually expresses hundreds of effector candidates, which may overwhelm host counter measures but make single effectors genetically dispensable (Spanu et al., 2010; Pedersen et al., 2012). Interestingly, 9o9 is not the only effector protein targeting activated RACB. The effector, *Bh* ROP-INTERACTIVE PEPTIDE 1 (ROPIP1), is an unconventional effector encoded by a transposable element and was previously found to interact with RACB resulting in the fragmentation of microtubules in the host cell, a process associated with susceptibility to fungal invasion (Huesmann *et al*., 2012, Nottensteiner *et al*., 2018). The finding that RACB is a potential target of two diverse effector proteins highlights the importance of RACB for the infection success of *Bh*. We currently can only speculate how 9o9 might manipulate RACB functions. In this context, it is worth noting that RACB function in cytoskeletal organization and cell polarity might be important for defense, too, because both processes have been shown to act in penetration resistance to diverse fungal pathogens (Schmelzer, 2002). It is possible that *Bh* takes advantage by abusing RACB function in cell polarity for redirecting transport processes towards material delivery for the formation of the haustorial complex, which depends on host-derived extrahaustorial matrix and membrane materials.

### PLC1 and SAC-like work in resistance against *Bh*

Besides 9o9, we also identified barley PLC1 and SAC-like as RACB-interactors in the RACB-CoIP experiment, and proteins were enriched in untreated or *Bh-*infected samples of the CoIP-LC-MS/MS-screening, respectively. During *Bh-*infection, *PLC1* and *SAC-like* were not differentially expressed over the first 24 h of host colonization (Fig. 3b, c). However, both proteins have roles in supporting resistance against powdery mildew infection, as the penetration frequency of *Bh* increased upon transient RNAi mediated silencing of gene expression (Fig. 4d). Both PLC1 and SAC-like are phosphoinositide-degrading enzymes, and their suppression likely contributes to the delimitation of phosphoinositide abundance. As phosphoinositides, such as PtdIns(4,5)P_2_, have been identified as susceptibility factors in different plant-pathogen interactions (Shimada *et al*., 2019, Qin *et al*., 2020), enhanced phosphoinositide abundance could explain the increased susceptibility to *Bh* observed in consequence of suppression of PLC1 and/or SAC-like in barley epidermis cells (Fig. 4d). It is interesting to see two enzymes of phosphoinositide-degradation to interact with RACB-CA in barley, adding to several other reported modes of limiting the abundance of phosphoinositides during plant defence, including the posttranslational inhibition of Arabidopsis PIP5K6 (Menzel *et al*., 2019) or the transcriptional activation of a gene encoding a phosphoinositide 5-phosphatase in potato in response to different PAMP treatments (Rausche *et al*., 2021). With regard to ROP signaling, canonical RACB-downstream interactors are often connected to powdery mildew susceptibility (Schultheiss *et al*., 2008, McCollum *et al*., 2020, Engelhardt *et al*., 2023). Other RACB-interacting proteins function in resistance against *Bh* and have been suggested to negatively regulate RACB-mediated signaling processes. For instance, MICROTUBULE ASSOCIATED ROP GTPase ACTIVATING PROTEIN1 likely inhibits RACB-signaling through its GTP-hydrolysing GAP-function, whereas ROP BINDING KINASE1 and the E3 ubiquitin ligase subunit SKP1L negatively regulate RACB protein abundance (Hoefle *et al*., 2011, Huesmann *et al*., 2012, Reiner *et al*., 2016). PLC1 and SAC-like might antagonize RACB-signaling via their predicted phospholipid-degrading activities, possibly limiting PM-association of RACB. Signaling pathways of RACB, PLC1 and SAC-like could converge on phosphoinositides because all three proteins can bind similar lipid species *in vitro* (Fig. 5). During *Bh*-attack, some anionic phospholipid species enrich at the site of fungal attack (Fig. 7) and thereby potentially recruit RACB in a similar manner as proposed for *At*ROP6 in *Arabidopsis* (Platre *et al*., 2019). Focal accumulation could subsequently support RACB-signaling processes that lead to *Bh*-susceptibility. However, enzymatic activity of PLC1 and SAC-like could degrade accumulating anionic phospholipids that would antagonize membrane polarization and related RACB signaling in susceptibility. In animals, activation of PLCs by RACB-like Rho-proteins has been shown (Jezyk *et al*., 2006, Rao *et al*., 2008, Li *et al*., 2009). In plants however, PLCs are activated by Ca^2+^ (Pokotylo *et al*., 2014), whereas little is known about the modes of regulation of SACs. PLC-proteins were shown to be heat stress-responsive and to govern drought and salt stress responses in different plant species (Liu *et al*., 2006, Wang *et al*., 2008, Georges *et al*., 2009, Tripathy *et al*., 2011). In biotic interactions, tomato PLC6 was shown to be important for defense against the fungi *Cladosporium fulvum* and *Verticillium dahliae*, and the bacterium *Pseudomonas syringae* (Vossen *et al*., 2010). In rice, gene expression of *OsPLC1*, the closest homolog of barley PLC1 (Fig. S4b), was shown to be stimulated by chemical and biological signals from the systemic acquired signaling pathway (Song and Goodman, 2002). In summary, these studies about different PLCs are consistent with a possible role of barley PLC1 in resistance to fungal pathogens. For proteins from the SAC-family, studies were conducted with regard to endomembrane trafficking and organelle morphology (Nováková *et al*., 2014, Mao and Tan, 2021).

### Dynamic asymmetric distribution of anionic phospholipids in barley cells during *Bh*-attack

Since we found phospholipid-binding abilities for RACB and its novel interaction partners, we also investigated the subcellular localization of anionic phospholipid markers in barley during *Bh*-attack. In *Bh*-infected barley epidermal cells, the plasma membrane-localized for PtdIns4P and PtdSer biomarkers were found enriched at the haustorial neck region, but were excluded from the plant extrahaustorial membrane (Fig. 7b, e, Fig. S4), whereas a biomarker for PtdIns(4,5)P_2_ was displaced from the PM during fungal attack (Fig. 7a, c). Some of our data match results from other plant-pathosystems, in which the subcellular distribution of different anionic phospholipid-species was investigated. For instance, in Arabidopsis, a PtdIns4P-biomarker was found at the haustorial neck region during infection by the oomycete, *Hyaloperonospora arabidopsidis* (*Hpa*) or by the powdery mildew fungus *Erysiphe cichoracearum* (*Ec*) (Shimada *et al*., 2019, Qin *et al*., 2020). In both pathosystems, the PtdIns4P-marker was also excluded from the extrahaustorial membrane. In Arabidopsis, markers for PtdIns(4,5) P_2_ appear enriched at the extrahaustorial membrane of *Hpa*, *Ec* and *Golovinomyces orontii* (*Go*) or the extra-invasive hyphal membrane of *Colletotrichum higginsianum* (Shimada *et al*., 2019, Qin *et al*., 2020), but were not clearly detected at comparable structures in barley epidermal cells. However, displacement of a PtdIns(4,5)P_2_-marker from infection structures was previously observed in potato plants infected by *Phytophthora infestans* (Rausche *et al*., 2021). Several anionic phospholipid-species have previously been linked to ROP-signaling. PtdSer is required for the auxin-induced partitioning of *At*ROP6 into PM-nanodomains during root gravitropism in Arabidopsis (Platre *et al*., 2019), whereas PtdIns(3,5)P_2_ might contribute to governing cell wall hardening in Arabidopsis root hairs together with the type II ROP, *At*ROP10 (Hirano *et al*., 2018). PtdIns(4,5) P_2_ mediates root hair positioning and polar growth in Arabidopsis together with the RACB-related type I ROPs, *At*ROP2 and *At*ROP6 (Jones *et al*., 2002, Kusano *et al*., 2008, Stanislas *et al*., 2015, Hirano *et al*., 2018), and it also regulates the actin cytoskeleton during *Nicotiana tabacum* (*Nt*) pollen tube growth via the type I ROP, *Nt*RAC5 (Kost *et al*., 1999, Fratini *et al*., 2021). In pollen tubes, PtdIns(4,5)P_2_ even stimulates the activation of ROPs, likely by interfering with the ROP interaction with GDP-dissociation inhibitors, which otherwise sequester ROPs in the cytoplasm and limit their activation at the PM (Kost, 2008, Ischebeck *et al*., 2011, Fratini *et al*., 2021). Together, the available information leads us to propose that anionic phospholipids serve as positional membrane signals for the PM recruitment of RACB and associated proteins to enable RACB-mediated susceptibility. This is supported by the lipid-dependent PM-association of RACB and RACB-supported plasma membrane association of its three canonical interactors RIC157, RIC171 and RIPb. When co-expressed during *Bh*-infection, RACB-CA and either of the three scaffold proteins co-localize at the haustorial neck region (Schultheiss *et al*., 2008, McCollum *et al*., 2020, Engelhardt *et al*., 2023), in a manner similar to what we observed for fluorescent reporters for PtdSer and particularly for PtdIns4P. Both RICs and RIPb contain polybasic stretches in their amino acid sequences, which possibly enables them to bind anionic phospholipids (Schultheiss *et al*., 2008, McCollum *et al*., 2020, Engelhardt *et al*., 2023).

The question arises how 9o9 might mechanistically support virulence by targeting RACB. Possibly an effector could influence RACB stability, GTP hydrolysis or the GDP for GTP-exchange of the ROP GTPase. Our protein complex modelling suggested that 9o9 bind to a set of surface amino acids in RACB that overlapped with RACB-PLC1 and RACB-SAC-like contact amino acids in the ROP effector loop that is also responsible for ROP regulators and downstream proteins (Fig. S5). Although modeling-based prediction has to be taken with care, one might hence speculate that *B. hordei* 9o9 competes with host proteins, which act in defense, for binding to RACB. Such competition could limit recruitment of antagonistic ROPGAPs or of PLC1 or SAC-like to the site of fungal attack. Since SAC-like is similar to SAC proteins that might act on PtdIns(3,5)P_2_, our observation that 9o9 supports accumulation of the PtdIns(3,5)P_2_-marker at the haustorial neck region could reflect displacement of barley SAC-like by fungal 9o9 leading to changes in the amount or subcellular distribution of PtdIns(3,5)P_2_. However, further experimental support is required to underpin this possible explanation. PtdIns(3,5)P_2_ is partially found on plant endosomes (Hirano *et al*. 2017; Noack and Jaillais, 2020). Possibly, 9o9 supports redirection of endosomes to the haustorial neck region or hinders early enough redirection during defense of fungal penetration attemps. In this context, it is interesting that host endomembrane trafficking is crucial for plant defense and target of other *B. hordei* effectors that suppress barley immunity (Nielsen, 2024; Liao *et al*. 2023; Sabelleck *et al*. 2025). In conclusion, we suggest that during *Bh*-infection anionic phospholipids enrich at infection sites and interact with susceptibility factors, such as RACB, which facilitates susceptibility towards *Bh*-infection and supports ingrowth of the fungal haustorium into the intact host cell. Simultaneously, RACB and anionic phospholipids may recruit phospholipid-metabolizing enzymes that function to spatially restrict phospholipid functions or membrane domains, and this might limit fungal infection success. Future studies may help to better understand the spatiotemporal order of lipid-and ROP-signaling events and the mechanistic detail of their interconnection.

## Experimental procedures

### Molecular cloning

Cloning of all constructs from this study was achieved by a combination of classical cloning, Gateway®-and GoldenGate-techniques. Please see Supplemental File 1 and Supplemental table 2 for a detailed description of all cloning processes and primers.

### Plant and fungal growth conditions

Wildtype barley (*Hordeum vulgare* L. subspecies *vulgare*) plants cultivar Golden Promise and transgenic barley lines were grown in a climate chamber (Conviron, Winnipeg, Canada) at 18 °C, 65% relative humidity and a cycle of 16 h of 150 µM s^-1^ m^-2^ light followed by 8 h dark. All experiments with transgenic plants used transgene-expressing individuals of generation T_2_ (see below; lines BG654 E02 and E12 for eGFP-RACB-CA, BG655 E01 and E10 for eGFP-RACB-CA-ΔCSIL and BG656 E01 and E06 for eGFP), with one exception: for the analysis of the *Bh* susceptibility of transgenic plants, transgene-expressing offspring of generation T_1_ were used, because the T_2_ plants were not ready at the time. The barley powdery mildew fungus *Blumeria hordei* isolate A6 (*Bh*) was cultivated on Golden Promise under the same conditions. All experiments used *Bh*-infected plants between 7-9 dpi as inoculum. *Nicotiana benthamiana* plants were sown on a mixture of 1 part vermiculite (1/3 mm, Raiffeisen Gartenbau, Köln, Germany) and 5 parts soil, then stratified for >2 days at 4 °C before being grown under long-day conditions (55% relative humidity, 16 h 150 µM s^-1^ m^-2^ light at 23 °C, 8 h dark at 21 °C).

### Generation and selection of transgenic barley lines

The transgenic barley lines from this study were created similarly to those described in Weiss *et al*. (2022). Please see Supplemental File 1 for a detailed description of the method.

### Immunoprecipitation

First leaves were frozen in liquid N_2_ and homogenized using a mortar and pestle for protein IPs from transgenic barley plants. Proteins were extracted using 400 µl extraction buffer (10% (w/v) glycerol, 25 mM Tris pH 7.5, 1 mM EDTA, 150 mM NaCl, 10 mM DTT, 1 mM PMSF, 0.5% Nonidet P40 substitute, 1x protease inhibitor (Sigma-Aldrich, St. Louis, USA, P9599)) and a 30 min incubation on ice, followed by removing cell debris via 10 min centrifugation at 4 °C and 18000 g. For targeted CoIPs from transiently transformed *Nicotiana benthamiana* plants, 5 leaf discs (1.2 mm diameter) were taken with a biopsy puncher, frozen in liquid N_2_ and homogenized using a TissueLyser II (QIAGEN, Hilden, Germany). Proteins were extracted in 500 µl extraction buffer and cell debris was removed as described above. Immunoprecipitation from debris-cleared supernatants of barley or *Nicotiana benthamiana* samples was conducted according to standard methods. For one α-GFP IP, 10 µl GFP-Trap® magnetic agarose beads (gtma-10, Chromotek, Planegg, Germany) were used (20 µl/sample for interactome samples). Briefly, beads were equilibrated 3 times with 400 µl extraction buffer. After adding supernatants, samples were tumbled end-over-end for 1 h at 4 °C, then washed 5 times with washing buffer (10% (w/v) glycerol, 25 mM Tris pH 7.5, 1 mM EDTA, 150 mM NaCl, 1 mM PMSF, 1x protease inhibitor (P9599, Sigma-Aldrich, St. Louis, USA). Proteins were eluted in either 50 µl 2x SDS-loading dye (for routine IPs; 40% (w/v) glycerol, 200 mM Tris-HCl pH 6.8, 20% β-Mercaptoethanol, 8% (w/v) SDS, 0.02% (w/v) bromophenol blue;) or 100 µl 2x NuPAGE™ LDS Sample Buffer (for the epidermis interactome; NP0008, Thermo Fisher Scientific, Waltham, USA) and boiled at 95 °C for 20 min. In case that additional fractions apart from the IP eluate were needed, 12.5 μl of 4x SDS-loading dye were mixed and boiled with 37.5 μl of samples taken after protein extraction (“Input”), incubation with the tag-specific beads (“Unbound”) or the last washing step (“Wash”). All samples were analyzed via routine SDS-PAGE and α-GFP Western blotting or α-GFP/α-HA Western blotting for CoIP-analysis. Interactome samples were instead used for peptide identification via mass spectrometry. All antibodies can be found in Supplemental Table 4.

### Preparation of the barley epidermis RACB-interactome

This experiment used pools of transgene-expressing plants in generation T_2_ for eGFP-RACB-CA (lines BG654 E02 and E12), eGFP-RACB-CA-ΔCSIL (lines: BG655 E01 and E10) and eGFP (BG656 E01 and E06). For the epidermis interactome, three biological replicates were prepared per construct (eGFP, eGFP-RACB-CA or eGFP-RACB-CA-ΔCSIL) and condition (*Bh*-infected or mock). One biological replicate consisted of three technical replicates. For one technical replicate of the “inoculated” condition, 21 primary leaves belonging to one construct were placed on 0.8% water-agar plates with the abaxial leaf side facing up. These plates were challenge-inoculated with 100-130 *Bh* spores/mm^2^ and put under normal growth conditions for 24 h. For the “mock” condition, the same approach without inoculum was used. After 24 hpi, abaxial epidermal peels were produced, pooled in one tube and stored in liquid nitrogen until homogenization. All interactome samples were homogenized using a TissueLyser II (QIAGEN, Hilden, Germany) and proteins were resuspended in 400 µl extraction buffer before being used in α-GFP immunoprecipitations.

### Mass spectrometry

Interactome IP eluates in LDS sample buffer were reduced with 10 mM dithiothreitol (DTT) for 1 h followed by alkylation with 55 mM chloroacetamide (CAA) for 30 min at room temperature. Samples were run into a 4-12% NuPAGE gel (Invitrogen, Carlsbad, USA) for approximately 1 cm. Samples from the same biological replicate and treatment condition were run together on one gel. In-gel digestion followed standard procedures with trypsin (Roche, Basel, Switzerland) and TEAB digestion buffer. Digested peptides were analyzed by liquid chromatography-coupled mass spectrometry (LC-MS/MS) analysis on Orbitrap mass spectrometers (Thermo Fisher Scientific, Waltham, USA) coupled on-line to a Dionex 3000 HPLC (Thermo Fisher Scientific, Waltham, USA). The liquid chromatography setup consisted of a 75 μm × 2 cm trap column packed in-house with Reprosil Pur ODS-3 5 μm particles (Dr. Maisch GmbH, Ammerbuch, Germany) and a 75 μm × 40 cm analytical column, packed with 3 μm particles of C18 Reprosil Gold 120 (Dr. Maisch GmbH, Ammerbuch, Germany).

Peptides were loaded onto the trap column using 0.1% FA in water. Measurements were performed on a Q Exactive HF-X (Thermo Fisher Scientific, Waltham, USA). Separation of the peptides was performed by using a linear gradient from 4% to 32% ACN with 5% DMSO, 0.1% formic acid in water over 20 min followed by a washing step (30 min total method length) at a flow rate of 300 nl/min and a column temperature of 50 °C. The instrument was operated in data-dependent mode, automatically switching between MS and MS2 scans. Full-scan mass spectra (m/z 360-1300) were acquired in profile mode with 60,000 resolution, an automatic gain control (AGC) target value of 3e6 and 45 ms maximum injection time. For the top 12 precursor ions, high resolution MS2 scans were performed using HCD fragmentation with 26% normalized collision energy, 15,000 resolution, an AGC target value of 2e5, 25 ms maximum injection time and 1.3 m/z isolation width in centroid mode. The minimum AGC target value was set to 2.2e3 with a dynamic exclusion of 20s. Peptide identification and quantification was performed with MaxQuant (Cox and Mann, 2008) using standard settings (V1.5.8.3). Raw files were searched against a barley database (Morex V2, 160517_Hv_IBSC_PGSB_r1_proteins_HighConf_REPR_annotation.fasta; IPK Gatersleben, Mascher *et al*. (2017) and common contaminants .The recombinant protein sequences and a barley powdery mildew reference database (*Blumeria hordei* isolate DH14, uniprot-proteome_UP000015441, Frantzeskakis *et al*. (2018)) were added to the search space. Carbamidomethylated cysteine was set as fixed modification and oxidation of methionine, N-terminal protein acetylation, phosphorylation of serine, threonine or tyrosine and GlyGly modification of lysine as variable modifications. Trypsin/P was specified as the proteolytic enzyme, with up to two missed cleavage sites allowed. The match between run function was enabled but restricted to either control or barley powdery mildew-treated samples. Results were filtered to 1% PSM, protein and Site FDR. Resulting data from MaxQuant was imported into Perseus V1.5.5.3 (Tyanova *et al*., 2016) for statistical analysis. Significant enrichment of peptides was calculated using two-tailed Student’s *t-*tests against an α of 0.05. Data was then imported into Microsoft Excel 2016 (Microsoft, Redmond, USA) for identification of novel eGFP-RACB-CA interaction partners. The considered criteria were: statistically significant enrichment or exclusive presence in eGFP-RACB-CA samples, degree of enrichment and number of unique peptides. The mass spectrometry proteomics data have been deposited to the ProteomeXchange Consortium via the PRIDE (PMID: 34723319) partner repository with the dataset identifier PXD056811.

### Transient transformation of *Nicotiana benthamiana* plants

*Nicotiana benthamiana* plants were transformed via *Agrobacterium tumefaciens* (strain GV3101 pMP90; Lazo *et al*. (1991)) with a protocol adapted from Yang *et al*. (2000). Please see Supplemental File 1 for a complete description.

### Transient transformation of barley

Epidermal cells of wild type barley *cv.* Golden Promise leaves were transiently transformed via particle bombardment with a protocol adapted from (Schweizer *et al*., 1999). See Supplemental File 1 for a comprehensive description.

### Analysis of *Bh* penetration efficiency

The *Bh* susceptibility assessment of transgenic plants was conducted in lines BG654 E2 for eGFP-RACB-CA, BG655 E1 for eGFP-RACB-CA-ΔCSIL and BG656 E1 for eGFP as described in Weiss *et al*. (2022). All plants were selected, transgene-expressing offspring in generation T_1_. *Bh* susceptibility of transiently transformed barley plants was performed as described in Engelhardt *et al*. (2023). For a comprehensive description, see Supplemental File 1.

### Confocal laser scanning microscopy

For subcellular localization experiments, barley leaf epidermis was transiently transformed as described above and imaged at 24 h after transformation (hat). In case of *Bh* infection, transformed barley leaves were inoculated with 100-130 *Bh* spores/mm^2^ at 8 hat and imaged at 16-20 hpi unless indicated otherwise. All subcellular localization experiments were performed with a Leica (Wetzlar, Germany) TCS SP5 mounted on a DM6000 stage. All images were taken with a HCX PL APO lambda blue 20.0x0.7 IMM UV objective (Leica, Wetzlar, Germany). CFP was excited with a 458 nm Argon laser line and detected between 463-485 nm; GFP was excited with a 488 nm Argon laser line and detected between 500-550 nm; mCitrine was excited with a 514 nm Argon laser line and detected between 525-550 nm; mCherry was excited with a 561 nm DPSS diode laser and detected between 570-620 nm. Highly fluorescent samples were imaged with a PMT, whereas samples with low fluorescence were analyzed with HyDs (both Leica, Wetzlar, Germany). For simultaneous imaging of multiple fluorophores, the sequential scan mode “between lines” was used. A line average of 3 was used for all scans. All images were captured as Z-stacks of single XY-optical sections with the Z-step size of each experiment indicated in figure legends. Image analysis was performed with Leica LAS X V3.5.1 (Leica, Wetzlar, Germany). λ-scanning was performed by selecting an area of interest that was in focus and exhibited strong fluorescence. For measurements, the Leica TCS SP5 was run in xyλ-mode and the fluorescence emission was detected in 5 nm bins in a range of 500-760 nm (GFP) or 525-760 nm (mCitrine) using HyDs. Fluorophores were excited with a 488 nm (GFP) or 514 nm (mCitrine) Argon laser line. After scanning, λ emission spectra were analyzed in selected regions-of-interest using Leica LAS X V3.5.1 (Leica, Wetzlar, Germany). Results were exported as normalized mean fluorescence intensities and plotted in GraphPad Prism V8.0 (GraphPad Software, San Diego, USA).

### FRET-FLIM measurements

FRET-FLIM measurements were performed with an FCS/FLIM-FRET/rapidFLIM upgrade kit (Picoquant, Berlin, Germany) used in tandem with an Olympus (Tokyo, Japan) FV3000 mounted on an IX63 stand. This method was adapted from (Weidtkamp-Peters and Stahl, 2017). Please see Supplemental File 1 for a detailed description of the technique.

### Yeast two-hybrid assay

The yeast two-hybrid assays was adapted from (Engelhardt *et al*., 2023). Please see Supplemental File 1 for a detailed description of the cloning.

### RT-qPCR

Gene expression of *9o9*, *PLC1* and *SAC-like* was measured using RT-qPCR in mock-treated or *Bh-*infected barley epidermal peels at the indicated timepoints. As housekeeping gene, *ubiquitin-conjugating enzyme 2* (*UBC2*) was used for barley, while β*-tubulin 2* (β*-TUB2*) was used for *Bh* (Schnepf *et al*., 2018). Please see Supplemental File 1 for a detailed description of the method.

### Production and purification of recombinant proteins

For *in vitro* lipid-binding assays, proteins were produced in *E. coli* (strain Rosetta 2). For that, chemically competent bacteria were transformed with pGEX-and pMAL-plasmids containing the constructs-of-interest (see molecular cloning). One colony per construct was taken to inoculate 30 ml 2YT (10 g/l yeast extract, 16 g/l tryptone, 5 g/l NaCl, 2 g/l D-glucose) starter culture that was grown overnight at 18 °C in a shaker. Next day, 300 ml of fresh 2YT medium were inoculated with 3 ml of starter culture and grown at 37 °C under constant shaking until an oD_600nm_ of 0.6-0.8 was reached. Bacterial growth was stopped by a 30 min incubation on ice, after which protein expression was started through addition of IPTG (final concentration: 0.1 mM). After induction, cultures were incubated for 18-20 h at 18 °C under constant shaking. All cultures were harvested in 50 ml aliquots via centrifugation at 3000 g and 4 °C. Cell pellets were frozen in liquid N_2_ and stored at -80 °C until use. Cell pellets were resuspended in 3 ml extraction buffer and incubated on ice for 30-45 min with occasional mixing. The extraction buffer for MBP-tagged proteins was: 20 mM Tris-HCl pH 7.4, 200 mM NaCl, 1 mM EDTA, 1x SigmaFast protease inhibitor cocktail without EDTA (Sigma-Aldrich, St. Louis, USA), 1 mM DTT, 2 mg/ml lysozyme. For GST-tagged proteins, the extraction buffer was: 50 mM Tris-HCl pH 7.4, 150 mM NaCl, 1x SigmaFast protease inhibitor cocktail without EDTA (Sigma-Aldrich, St. Louis, USA), 1 mM DTT, 2 mg/ml lysozyme. After resuspension, cells were lysed through sonication, delivering 4000 J in 2 s pulses using a Vibra-Cell^TM^ 72442 with Bransonic B12 Ultrasonics Sonifier (both from Branson Ultrasonics, Brookfield, USA). Cell lysates were centrifuged at 20000 g for 15 min at 4 °C to separate supernatant and pellet fractions. The protein extracts in the supernatants were finally used for protein purification via affinity chromatography. MBP-fusion proteins were enriched via amylose resin (NEB, Ipswich, USA), while GST-fusion proteins were enriched Pierce^TM^ glutathione agarose resin (Thermo Fisher Scientific, Waltham USA). Both resins were packed into Pierce^TM^ centrifugation columns (Thermo Fisher Scientific, Waltham, USA) before use. To start, both columns were washed once with 3 ml ddH_2_O and twice with either 3 ml MBP equilibration buffer (20 mM Tris-HCl pH 7.4, 200 mM NaCl, 1 mM EDTA, 1 mM DTT) or 3 ml GST equilibration buffer (50 mM Tris-HCl pH 7.4, 150 mM NaCl, 1 mM DTT). Between washing steps, columns were centrifuged for 1 min at 400 g and 4 °C. Afterwards, raw protein extracts were added onto the columns and incubated for 1 h at room temperature in an overhead tumbler. Followingly, each column was washed three times with its respective equilibration buffer from above. Lastly, fusion proteins were eluted in either MBP elution buffer (10 mM maltose, 20 mM Tris-HCl pH 7.4, 200 mM NaCl, 1 mM EDTA, 1 mM DTT) or GST elution buffer (50 mM glutathion, 50 mM Tris-HCl pH 7.4, 150 mM NaCl, 1 mM DTT). Proteins were stored at -80 °C until use.

### Protein-lipid-overlay experiments

Commercially available membranes with pre-spotted lipids (PIP Strips, Echelon Biosciences Inc., MoBiTec GmbH, Göttingen, Germany) were first blocked with 3% (w/v) skimmed milk powder in TBS (50 mM Tris-HCl pH 7.5, 150 mM NaCl) for 30 min. Subsequently, membranes were incubated overnight with 0.5 µg/ml purified proteins in 3% skimmed milk in TBS at 4 °C and gentle shaking. Next day, membranes were washed three times with TBS and incubated with primary antibodies in 3% skimmed milk in TBS for 1 h at room temperature with gentle shaking. Membranes were again washed three times with TBS, before being incubated with secondary antibodies in 3% skimmed milk in TBS for 1 h at room temperature with gentle shaking. Afterwards, membranes were washed twice with TBS and once with AP-buffer (100 mM Tris-HCl pH 9.5, 100 mM NaCl, 5 mM MgCl_2_). Proteins were detected using 0.175 mg/ml BCIP (5-bromo-4-chloro-3-indolyl phosphate di-sodium salt, Roth, Karlsruhe, Germany) and 0.338 mg/ml NBT (nitro blue tetrazolium chloride, Roth, Karlsruhe, Germany) in AP-buffer. Reactions were stopped by washing with ddH_2_O, when sufficient staining was attained. All antibodies can be found in Supplemental Table 4.

### Bioinformatic analyses

All bioinformatics analyses are described in Supplemental File 1.

## Author contributions

LSW and RH designed the study concept. All authors designed experiments. LSW, CB, MB, JM, MH, GH performed the experiments. RH, BK, JK, JH, SE supervised the work. RH, JK, JH and BK provided resources. LSW and RH wrote the manuscript draft. All authors critically read and revised the manuscript and approved the final version.

## Acknowledgements

We thank Yvon Jaillais (ENS Lyon) for providing the pB7m34GW-pUBQ10::Citrine-C2^LACT^ plasmid and Masa Sato (Kyoto Prefectural University) for sending the pGWB501-UBQ10::mCitrine-2xML1N plasmid. We acknowledge Cornelia Marthe for technical assistance in generating the transgenic barley lines, and thank Johanna Hofer and Vanessa Weiß for outstanding technical assistance in wet lab work at TU Munich. We thank Marina Hijano Moreno and Tina Reiner (TU Munich) for minor experimental contributions. We acknowledge the TUM Center for Advanced Light Microscopy (CALM) at the TUM School of Life Sciences Weihenstephan for providing their FRET-FLIM equipment and training and TUM Plant Technology Center for support in propagation of transgenic barley seed. This study was supported by the German Research foundation with grants within collaborative research centers SFB624 (grants to R.H. and B.K.) and HU886/12 (to R.H.).

## Conflict of interests

The authors declare no conflict of interest.

## Supporting information

Additional Supporting Information may be found in the online version of this article.

Figure S1: RACB-CA epidermis interactome screening.

Figure S2: Annotation of functional domains and catalytic amino acids in PLC1 and SAC-like.

Figure S3: Homologs of 9o9, PLC1 and SAC-like in barley, rice, Arabidopsis and Bh. Figure S1

Figure S4: Zoom in magnification of phospholipid marker accumulation at sites of penetration by *B. hordei*.

Figure S5: Interaction of 9o9 with RACB *in vivo* and of 9o9, PLC1 and SAC-like with RACB in silico.

Supplemental Table S1: Full dataset from the barley eGFP-RACB-CA epidermis interactome.

Supplemental Table S2: Primers.

Supplemental Table S3: Genes.

Supplemental Table S4: Antibodies.

Supplemental File 1: Additional Experimental Procedures and references.

## Notes

### Competing Interest Statement

The authors have declared no competing interest.

### Summary of Updates

revised manuscript resubmitted for final acceptance

## References

1. Akamatsu, A., Wong, H.L., Fujiwara, M., Okuda, J., Nishide, K., Uno, K., Imai, K., Umemura, K., Kawasaki, T., Kawano, Y. and Shimamoto, K. (2013) An OsCEBiP/OsCERK1-OsRacGEF1-OsRac1 module is an essential early component of chitin-induced rice immunity. Cell Host Microbe, 13, 465–476.

2. Bak, G., Lee, E.J., Lee, Y., Kato, M., Segami, S., Sze, H., Maeshima, M., Hwang, J.U. and Lee, Y. (2013) Rapid structural changes and acidification of guard cell vacuoles during stomatal closure require phosphatidylinositol 3,5-bisphosphate. Plant Cell, 25, 2202–2216.

3. Berken, A. (2006) ROPs in the spotlight of plant signal transduction. Cellular and Molecular Life Sciences, 63, 2446–2459.

4. Caldelari, D., Sternberg, H., Rodrıǵuez-Concepción, M., Gruissem, W. and Yalovsky, S. (2001) Efficient Prenylation by a Plant Geranylgeranyltransferase-I Requires a Functional CaaL Box Motif and a Proximal Polybasic Domain. Plant Physiology, 126, 1416–1429.

5. Choi, S., Prokchorchik, M., Lee, H., Gupta, R., Lee, Y., Chung, E.H., Cho, B., Kim, M.S., Kim, S.T. and Sohn, K.H. (2021) Direct acetylation of a conserved threonine of RIN4 by the bacterial effector HopZ5 or AvrBsT activates RPM1-dependent immunity in Arabidopsis. Molecular plant, 14, 1951–1960.

6. Choi, Y., Lee, Y., Kim, S.Y., Lee, Y. and HWANG, J.U. (2013) Arabidopsis ROP-interactive CRIB motif-containing protein 1 (RIC1) positively regulates auxin signalling and negatively regulates abscisic acid (ABA) signalling during root development. Plant, Cell & Environment, 36, 945–955.

7. Cox, J. and Mann, M. (2008) MaxQuant enables high peptide identification rates, individualized ppb-range mass accuracies and proteome-wide protein quantification. Nature biotechnology, 26, 1367–1372.

8. Engelhardt, S., Trutzenberg, A., Kopischke, M., Probst, K., McCollum, C., Hofer, J. and Hückelhoven, R. (2023) Barley RIC157, a potential RACB scaffold protein, is involved in susceptibility to powdery mildew. Plant Molecular Biology, 111, 329–344.

9. Frantzeskakis, L., Kracher, B., Kusch, S., Yoshikawa-Maekawa, M., Bauer, S., Pedersen, C., Spanu, P.D., Maekawa, T., Schulze-Lefert, P. and Panstruga, R. (2018) Signatures of host specialization and a recent transposable element burst in the dynamic one-speed genome of the fungal barley powdery mildew pathogen. BMC Genomics, 19, 381.

10. Fratini, M., Krishnamoorthy, P., Stenzel, I., Riechmann, M., Matzner, M., Bacia, K., Heilmann, M. and Heilmann, I. (2021) Plasma membrane nano-organization specifies phosphoinositide effects on Rho-GTPases and actin dynamics in tobacco pollen tubes. Plant Cell, 33, 642–670.

11. Fu, Y., Xu, T., Zhu, L., Wen, M. and Yang, Z. (2009) A ROP GTPase Signaling Pathway Controls Cortical Microtubule Ordering and Cell Expansion in Arabidopsis. Current Biology, 19, 1827–1832.

12. Georges, F., Das, S., Ray, H., Bock, C., Nokhrina, K., Kolla, V.A. and Keller, W. (2009) Over-expression of Brassica napus phosphatidylinositol-phospholipase C2 in canola induces significant changes in gene expression and phytohormone distribution patterns, enhances drought tolerance and promotes early flowering and maturation. Plant Cell Environ, 32, 1664–1681.

13. Gerth, K., Lin, F., Menzel, W., Krishnamoorthy, P., Stenzel, I., Heilmann, M. and Heilmann, I. (2017) Guilt by Association: A Phenotype-Based View of the Plant Phosphoinositide Network. Annu Rev Plant Biol, 68, 349–374.

14. Gu, Y., Fu, Y., Dowd, P., Li, S., Vernoud, V., Gilroy, S. and Yang, Z. (2005) A Rho family GTPase controls actin dynamics and tip growth via two counteracting downstream pathways in pollen tubes. J Cell Biol, 169, 127–138.

15. He, Q., Naqvi, S., McLellan, H., Boevink, P.C., Champouret, N., Hein, I. and Birch, P.R.J. (2018) Plant pathogen effector utilizes host susceptibility factor NRL1 to degrade the immune regulator SWAP70. Proc Natl Acad Sci U S A, 115, E7834–e7843.

16. Hirano, T., Konno, H., Takeda, S., Dolan, L., Kato, M., Aoyama, T., Higaki, T., Takigawa-Imamura, H. and Sato, M.H. (2018) PtdIns(3,5)P2 mediates root hair shank hardening in Arabidopsis. Nature Plants, 4, 888–897.

17. Hirano, T., Stecker, K., Munnik, T., Xu, H. and Sato, M.H. (2017) Visualization of Phosphatidylinositol 3,5-Bisphosphate Dynamics by a Tandem ML1N-Based Fluorescent Protein Probe in Arabidopsis. Plant Cell Physiol, 58, 1185–1195.

18. Hoefle, C., Huesmann, C., Schultheiss, H., Bornke, F., Hensel, G., Kumlehn, J. and Huckelhoven, R. (2011) A barley ROP GTPase ACTIVATING PROTEIN associates with microtubules and regulates entry of the barley powdery mildew fungus into leaf epidermal cells. Plant Cell, 23, 2422–2439.

19. Huesmann, C., Reiner, T., Hoefle, C., Preuss, J., Jurca, M.E., Domoki, M., Feher, A. and Huckelhoven, R. (2012) Barley ROP binding kinase1 is involved in microtubule organization and in basal penetration resistance to the barley powdery mildew fungus. Plant Physiol, 159, 311–320.

20. Ischebeck, T., Stenzel, I., Hempel, F., Jin, X., Mosblech, A. and Heilmann, I. (2011) Phosphatidylinositol-4,5-bisphosphate influences Nt-Rac5-mediated cell expansion in pollen tubes of Nicotiana tabacum. Plant J, 65, 453–468.

21. Jezyk, M.R., Snyder, J.T., Gershberg, S., Worthylake, D.K., Harden, T.K. and Sondek, J. (2006) Crystal structure of Rac1 bound to its effector phospholipase C-beta2. Nat Struct Mol Biol, 13, 1135–1140.

22. Jones, M.A., Shen, J.J., Fu, Y., Li, H., Yang, Z. and Grierson, C.S. (2002) The Arabidopsis Rop2 GTPase is a positive regulator of both root hair initiation and tip growth. Plant Cell, 14, 763–776.

23. Kost, B. (2008) Spatial control of Rho (Rac-Rop) signaling in tip-growing plant cells. Trends Cell Biol, 18, 119–127.

24. Kost, B., Lemichez, E., Spielhofer, P., Hong, Y., Tolias, K., Carpenter, C. and Chua, N.-H. (1999) Rac Homologues and Compartmentalized Phosphatidylinositol 4, 5-Bisphosphate Act in a Common Pathway to Regulate Polar Pollen Tube Growth. Journal of Cell Biology, 145, 317–330.

25. Kulich, I., Vogler, F., Bleckmann, A., Cyprys, P., Lindemeier, M., Fuchs, I., Krassini, L., Schubert, T., Steinbrenner, J., Beynon, J., Falter-Braun, P., Langst, G., Dresselhaus, T. and Sprunck, S. (2020) ARMADILLO REPEAT ONLY proteins confine Rho GTPase signalling to polar growth sites. Nat Plants, 6, 1275–1288.

26. Kusano, H., Testerink, C., Vermeer, J.E., Tsuge, T., Shimada, H., Oka, A., Munnik, T. and Aoyama, T. (2008) The Arabidopsis Phosphatidylinositol Phosphate 5-Kinase PIP5K3 is a key regulator of root hair tip growth. Plant Cell, 20, 367–380.

27. Kwaaitaal, M., Nielsen, M.E., Bohlenius, H. and Thordal-Christensen, H. (2017) The plant membrane surrounding powdery mildew haustoria shares properties with the endoplasmic reticulum membrane. J Exp Bot, 68, 5731–5743.

28. Lavy, M. and Yalovsky, S. (2006) Association of Arabidopsis type-II ROPs with the plasma membrane requires a conserved C-terminal sequence motif and a proximal polybasic domain. Plant Journal, 46, 934–947.

29. Lavy, M., Bloch, D., Hazak, O., Gutman, I., Poraty, L., Sorek, N., Sternberg, H. and Yalovsky, S. (2007) A Novel ROP/RAC effector links cell polarity, root-meristem maintenance, and vesicle trafficking. Curr Biol, 17, 947–952.

30. Lazo, G.R., Stein, P.A. and Ludwig, R.A. (1991) A DNA transformation–competent Arabidopsis genomic library in Agrobacterium. Bio/technology, 9, 963–967.

31. Liao, W., Nielsen, M.E., Pedersen, C., Xie, W. and Thordal-Christensen, H. (2023) Barley endosomal MONENSIN SENSITIVITY1 is a target of the powdery mildew effector CSEP0162 and plays a role in plant immunity. J Exp Bot, 74, 118–129.

32. Li, S., Gu, Y., Yan, A., Lord, E. and Yang, Z.B. (2008) RIP1 (ROP Interactive Partner 1)/ICR1 marks pollen germination sites and may act in the ROP1 pathway in the control of polarized pollen growth. Molecular plant, 1, 1021–1035.

33. Li, S., Wang, Q., Wang, Y., Chen, X. and Wang, Z. (2009) PLC-gamma1 and Rac1 coregulate EGF-induced cytoskeleton remodeling and cell migration. Mol Endocrinol, 23, 901–913.

34. Lin, D., Cao, L., Zhou, Z., Zhu, L., Ehrhardt, D., Yang, Z. and Fu, Y. (2013) Rho GTPase signaling activates microtubule severing to promote microtubule ordering in Arabidopsis. Current Biology, 23, 290–297.

35. Liu, H.-T., Huang, W.-D., Pan, Q.-H., Weng, F.-H., Zhan, J.-C., Liu, Y., Wan, S.-B. and Liu, Y.-Y. (2006) Contributions of PIP2-specific-phospholipase C and free salicylic acid to heat acclimation-induced thermotolerance in pea leaves. Journal of plant physiology, 163, 405–416.

36. Lück, S., Kreszies, T., Strickert, M., Schweizer, P., Kuhlmann, M. and Douchkov, D. (2019) siRNA-Finder (si-Fi) software for RNAi-target design and off-target prediction. Frontiers in Plant Science, 10, 1023.

37. Mackey, D., Holt III, B.F., Wiig, A. and Dangl, J.L. (2002) RIN4 interacts with Pseudomonas syringae type III effector molecules and is required for RPM1-mediated resistance in Arabidopsis. Cell, 108, 743–754.

38. Mao, Y. and Tan, S. (2021) Functions and Mechanisms of SAC Phosphoinositide Phosphatases in Plants. Front Plant Sci, 12, 803635.

39. Mascher, M., Gundlach, H., Himmelbach, A., Beier, S., Twardziok, S.O., Wicker, T., Radchuk, V., Dockter, C., Hedley, P.E., Russell, J., Bayer, M., Ramsay, L., Liu, H., Haberer, G., Zhang, X.Q., Zhang, Q., Barrero, R.A., Li, L., Taudien, S., Groth, M., Felder, M., Hastie, A., Simkova, H., Stankova, H., Vrana, J., Chan, S., Munoz-Amatriain, M., Ounit, R., Wanamaker, S., Bolser, D., Colmsee, C., Schmutzer, T., Aliyeva-Schnorr, L., Grasso, S., Tanskanen, J., Chailyan, A., Sampath, D., Heavens, D., Clissold, L., Cao, S., Chapman, B., Dai, F., Han, Y., Li, H., Li, X., Lin, C., McCooke, J.K., Tan, C., Wang, P., Wang, S., Yin, S., Zhou, G., Poland, J.A., Bellgard, M.I., Borisjuk, L., Houben, A., Dolezel, J., Ayling, S., Lonardi, S., Kersey, P., Langridge, P., Muehlbauer, G.J., Clark, M.D., Caccamo, M., Schulman, A.H., Mayer, K.F.X., Platzer, M., Close, T.J., Scholz, U., Hansson, M., Zhang, G., Braumann, I., Spannagl, M., Li, C., Waugh, R. and Stein, N. (2017) A chromosome conformation capture ordered sequence of the barley genome. Nature, 544, 427–433.

40. McCollum, C., Engelhardt, S., Weiss, L. and Huckelhoven, R. (2020) ROP INTERACTIVE PARTNER b Interacts with RACB and Supports Fungal Penetration into Barley Epidermal Cells. Plant Physiol, 184, 823–836.

41. Menzel, W., Stenzel, I., Helbig, L.M., Krishnamoorthy, P., Neumann, S., Eschen-Lippold, L., Heilmann, M., Lee, J. and Heilmann, I. (2019) A PAMP-triggered MAPK cascade inhibits phosphatidylinositol 4,5-bisphosphate production by PIP5K6 in Arabidopsis thaliana. New Phytol, 224, 833–847.

42. Mueller-Roeber, B. and Pical, C. (2002) Inositol phospholipid metabolism in Arabidopsis. Characterized and putative isoforms of inositol phospholipid kinase and phosphoinositide-specific phospholipase C. Plant Physiol, 130, 22–46.

43. Caldelari, D., Sternberg, H., Rodrıǵuez-Concepción, M., Gruissem, W. and Yalovsky, S. (2001) Efficient Prenylation by a Plant Geranylgeranyltransferase-I Requires a Functional CaaL Box Motif and a Proximal Polybasic Domain. Plant Physiology, 126, 1416–1429.

44. Heo, W.D., Inoue, T., Park, W.S., Kim, M.L., Park, B.O., Wandless, T.J. and Meyer, T. (2006) PI(3,4,5)P3 and PI(4,5)P2 Lipids Target Proteins with Polybasic Clusters to the Plasma Membrane. Science, 314, 1458–1461.

45. Lavy, M. and Yalovsky, S. (2006) Association of Arabidopsis type-II ROPs with the plasma membrane requires a conserved C-terminal sequence motif and a proximal polybasic domain. Plant Journal, 46, 934–947.

46. Liao, W., Nielsen, M.E., Pedersen, C., Xie, W. and Thordal-Christensen, H. (2023) Barley endosomal MONENSIN SENSITIVITY1 is a target of the powdery mildew effector CSEP0162 and plays a role in plant immunity. J Exp Bot, 74, 118–129.

47. Nielsen, M.E. (2024) Vesicle trafficking pathways in defence-related cell wall modifications: papillae and encasements. J Exp Bot, 75, 3700–3712.

48. Noack, L.C. and Jaillais, Y. (2020) Functions of Anionic Lipids in Plants. Annual Review of Plant Biology, 71, 71–102.

49. Nottensteiner, M., Zechmann, B., McCollum, C. and Huckelhoven, R. (2018) A barley powdery mildew fungus non-autonomous retrotransposon encodes a peptide that supports penetration success on barley. J Exp Bot, 69, 3745–3758.

50. Nováková, P., Hirsch, S., Feraru, E., Tejos, R., van Wijk, R., Viaene, T., Heilmann, M., Lerche, J., De Rycke, R., Feraru, M.I., Grones, P., Van Montagu, M., Heilmann, I., Munnik, T. and Friml, J. (2014) SAC phosphoinositide phosphatases at the tonoplast mediate vacuolar function in Arabidopsis. Proc Natl Acad Sci U S A, 111, 2818–2823.

51. Platre, M.P., Bayle, V., Armengot, L., Bareille, J., Marquès-Bueno, M.D.M., Creff, A., Maneta-Peyret, L., Fiche, J.-B., Nollmann, M. and Miège, C. (2019) Developmental control of plant Rho GTPase nano-organization by the lipid phosphatidylserine. Science, 364, 57–62.

52. Platre, M.P., Noack, L.C., Doumane, M., Bayle, V., Simon, M.L.A., Maneta-Peyret, L., Fouillen, L., Stanislas, T., Armengot, L., Pejchar, P., Caillaud, M.C., Potocky, M., Copic, A., Moreau, P. and Jaillais, Y. (2018) A Combinatorial Lipid Code Shapes the Electrostatic Landscape of Plant Endomembranes. Dev Cell, 45, 465–480 e411.

53. Pedersen, C., van Themaat, E.V.L., McGuffin, L., Abbott, J., Burgis, T., Barton, G., Bindschedler, L., Lu, X., Maekawa, T., WeSZling, R., Cramer, R., Thordal-Christensen, H., Panstruga, R. and Spanu, P. (2012) Structure and evolution of barley powdery mildew effector candidates. BMC Genomics, 13, 694.

54. Pokotylo, I., Kolesnikov, Y., Kravets, V., Zachowski, A. and Ruelland, E. (2014) Plant phosphoinositide-dependent phospholipases C: variations around a canonical theme. Biochimie, 96, 144–157.

55. Qin, L., Zhou, Z., Li, Q., Zhai, C., Liu, L., Quilichini, T.D., Gao, P., Kessler, S.A., Jaillais, Y., Datla, R., Peng, G., Xiang, D. and Wei, Y. (2020) Specific Recruitment of Phosphoinositide Species to the Plant-Pathogen Interfacial Membrane Underlies Arabidopsis Susceptibility to Fungal Infection. Plant Cell, 32, 1665–1688.

56. Rao, J.N., Liu, S.V., Zou, T., Liu, L., Xiao, L., Zhang, X., Bellavance, E., Yuan, J.X. and Wang, J.Y. (2008) Rac1 promotes intestinal epithelial restitution by increasing Ca2+ influx through interaction with phospholipase C-(gamma)1 after wounding. Am J Physiol Cell Physiol, 295, C1499–1509.

57. Rapazote-Flores, P., Bayer, M., Milne, L., Mayer, C.D., Fuller, J., Guo, W., Hedley, P.E., Morris, J., Halpin, C., Kam, J., McKim, S.M., Zwirek, M., Casao, M.C., Barakate, A., Schreiber, M., Stephen, G., Zhang, R., Brown, J.W.S., Waugh, R. and Simpson, C.G. (2019) BaRTv1.0: an improved barley reference transcript dataset to determine accurate changes in the barley transcriptome using RNA-seq. BMC Genomics, 20, 968.

58. Rausche, J., Stenzel, I., Stauder, R., Fratini, M., Trujillo, M., Heilmann, I. and Rosahl, S. (2021) A phosphoinositide 5-phosphatase from Solanum tuberosum is activated by PAMP-treatment and may antagonize phosphatidylinositol 4,5-bisphosphate at Phytophthora infestans infection sites. New Phytol, 229, 469–487.

59. Reiner, T., Hoefle, C. and Huckelhoven, R. (2016) A barley SKP1-like protein controls abundance of the susceptibility factor RACB and influences the interaction of barley with the barley powdery mildew fungus. Mol Plant Pathol, 17, 184–195.

60. Sabelleck, B., Deb, S., Levecque, S.C.J., Freh, M., Reinstädler, A., Spanu, P.D., Thordal-Christensen, H. and Panstruga, R. (2025) A powdery mildew core effector protein targets the host endosome tethering complexes HOPS and CORVET in barley. Plant Physiology, 197, kiaf067.

61. Saur, I.M.L. and Hückelhoven, R. (2021) Recognition and defence of plant-infecting fungal pathogens. Journal of plant physiology, 256, 153324.

62. Schmelzer, E. (2002) Cell polarization, a crucial process in fungal defence. Trends Plant Sci, 7, 411–415.

63. Schnepf, V., Vlot, A.C., Kugler, K. and Hückelhoven, R. (2018) Barley susceptibility factor RACB modulates transcript levels of signalling protein genes in compatible interaction with Blumeria graminis f.sp. hordei. Mol Plant Pathol, 19, 393–404.

64. Schultheiss, H., Dechert, C., Kogel, K.H. and Huckelhoven, R. (2002) A small GTP-binding host protein is required for entry of powdery mildew fungus into epidermal cells of barley. Plant Physiol, 128, 1447–1454.

65. Schultheiss, H., Dechert, C., Kogel, K.H. and Huckelhoven, R. (2003) Functional analysis of barley RAC/ROP G-protein family members in susceptibility to the powdery mildew fungus. Plant J, 36, 589–601.

66. Schultheiss, H., Preuss, J., Pircher, T., Eichmann, R. and Huckelhoven, R. (2008) Barley RIC171 interacts with RACB in planta and supports entry of the powdery mildew fungus. Cell Microbiol, 10, 1815–1826.

67. Schweizer, P., Christoffel, A. and Dudler, R. (1999) Transient expression of members of the germin-like gene family in epidermal cells of wheat confers disease resistance. The Plant Journal, 20, 541–552.

68. Sherwood, J.E. and Somerville, S.C. (1990) Sequence of the Erysiphe graminis f. sp. hordei gene encoding beta-tubulin. Nucleic acids research, 18, 1052.

69. Shimada, T.L., Betsuyaku, S., Inada, N., Ebine, K., Fujimoto, M., Uemura, T., Takano, Y., Fukuda, H., Nakano, A. and Ueda, T. (2019) Enrichment of Phosphatidylinositol 4,5-Bisphosphate in the Extra-Invasive Hyphal Membrane Promotes Colletotrichum Infection of Arabidopsis thaliana. Plant Cell Physiol, 60, 1514–1524.

70. Simon, M.L., Platre, M.P., Assil, S., van Wijk, R., Chen, W.Y., Chory, J., Dreux, M., Munnik, T. and Jaillais, Y. (2014) A multi-colour/multi-affinity marker set to visualize phosphoinositide dynamics in Arabidopsis. Plant J, 77, 322–337.

71. Spanu, P.D., Abbott, J.C., Amselem, J., Burgis, T.A., Soanes, D.M., Stuber, K., Ver Loren van Themaat, E., Brown, J.K., Butcher, S.A., Gurr, S.J., Lebrun, M.H., Ridout, C.J., Schulze-Lefert, P., Talbot, N.J., Ahmadinejad, N., Ametz, C., Barton, G.R., Benjdia, M., Bidzinski, P., Bindschedler, L.V., Both, M., Brewer, M.T., Cadle-Davidson, L., Cadle-Davidson, M.M., Collemare, J., Cramer, R., Frenkel, O., Godfrey, D., Harriman, J., Hoede, C., King, B.C., Klages, S., Kleemann, J., Knoll, D., Koti, P.S., Kreplak, J., Lopez-Ruiz, F.J., Lu, X., Maekawa, T., Mahanil, S., Micali, C., Milgroom, M.G., Montana, G., Noir, S., O’Connell, R.J., Oberhaensli, S., Parlange, F., Pedersen, C., Quesneville, H., Reinhardt, R., Rott, M., Sacristan, S., Schmidt, S.M., Schon, M., Skamnioti, P., Sommer, H., Stephens, A., Takahara, H., Thordal-Christensen, H., Vigouroux, M., Wessling, R., Wicker, T. and Panstruga, R. (2010) Genome expansion and gene loss in powdery mildew fungi reveal tradeoffs in extreme parasitism. Science, 330, 1543–1546.

72. Song, F. and Goodman, R.M. (2002) Molecular cloning and characterization of a rice phosphoinositide-specific phospholipase C gene, OsPI-PLC1, that is activated in systemic acquired resistance. Physiological and molecular plant pathology, 61, 31–40.

73. Sorek, N., Poraty, L., Sternberg, H., Buriakovsky, E., Bar, E., Lewinsohn, E. and Yalovsky, S. (2017) Corrected and Republished from: Activation Status-Coupled Transient S-Acylation Determines Membrane Partitioning of a Plant Rho-Related GTPase. Mol Cell Biol, 37, e00333–17.

74. Stanislas, T., Huser, A., Barbosa, I.C., Kiefer, C.S., Brackmann, K., Pietra, S., Gustavsson, A., Zourelidou, M., Schwechheimer, C. and Grebe, M. (2015) Arabidopsis D6PK is a lipid domain-dependent mediator of root epidermal planar polarity. Nat Plants, 1, 15162.

75. Sternberg, H., Buriakovsky, E., Bloch, D., Gutman, O., Henis, Y.I. and Yalovsky, S. (2021) Formation of self-organizing functionally distinct Rho of plants domains involves a reduced mobile population. Plant Physiology, 187, 2485–2508.

76. Tripathy, M.K., Tyagi, W., Goswami, M., Kaul, T., Singla-Pareek, S.L., Deswal, R., Reddy, M.K. and Sopory, S.K. (2011) Characterization and Functional Validation of Tobacco PLC Delta for Abiotic Stress Tolerance. Plant Molecular Biology Reporter, 30, 488–497.

77. Trutzenberg, A., Engelhardt, S., Weiß, L. and Hückelhoven, R. (2022) Barley guanine nucleotide exchange factor Hv GEF14 is an activator of the susceptibility factor Hv RACB and supports host cell entry by Blumeria graminis f. sp. hordei. Molecular plant pathology, 23, 1524–1537.

78. Tyanova, S., Temu, T., Sinitcyn, P., Carlson, A., Hein, M.Y., Geiger, T., Mann, M. and Cox, J. (2016) The Perseus computational platform for comprehensive analysis of (prote) omics data. Nature methods, 13, 731–740.

79. van Leeuwen, W., Vermeer, J.E., Gadella, T.W., Jr. and Munnik, T. (2007) Visualization of phosphatidylinositol 4,5-bisphosphate in the plasma membrane of suspension-cultured tobacco BY-2 cells and whole Arabidopsis seedlings. Plant J, 52, 1014–1026.

80. van Schie, C.C. and Takken, F.L. (2014) Susceptibility genes 101: how to be a good host. Annu Rev Phytopathol, 52, 551–581.

81. Vossen, J.H., Abd-El-Haliem, A., Fradin, E.F., van den Berg, G.C., Ekengren, S.K., Meijer, H.J., Seifi, A., Bai, Y., ten Have, A., Munnik, T., Thomma, B.P. and Joosten, M.H. (2010) Identification of tomato phosphatidylinositol-specific phospholipase-C (PI-PLC) family members and the role of PLC4 and PLC6 in HR and disease resistance. Plant J, 62, 224–239.

82. Wang, C.R., Yang, A.F., Yue, G.D., Gao, Q., Yin, H.Y. and Zhang, J.R. (2008) Enhanced expression of phospholipase C 1 (ZmPLC1) improves drought tolerance in transgenic maize. Planta, 227, 1127–1140.

83. Weidtkamp-Peters, S. and Stahl, Y. (2017) The Use of FRET/FLIM to Study Proteins Interacting with Plant Receptor Kinases. In Plant Receptor Kinases: Methods and Protocols (Aalen, R.B. ed. New York, NY: Springer New York, pp. 163–175.

84. Weiss, L., Gaelings, L., Reiner, T., Mergner, J., Kuster, B., Feher, A., Hensel, G., Gahrtz, M., Kumlehn, J., Engelhardt, S. and Huckelhoven, R. (2022) Posttranslational modification of the RHO of plants protein RACB by phosphorylation and cross-kingdom conserved ubiquitination. PLoS One, 17, e0258924.

85. Winge, P., Brembu, T., Kristensen, R. and Bones, A.M. (2000) Genetic structure and evolution of RAC-GTPases in Arabidopsis thaliana. Genetics, 156, 1959–1971.

86. Wu, G., Gu, Y., Li, S. and Yang, Z. (2001) A genome-wide analysis of Arabidopsis Rop-interactive CRIB motif–containing proteins that act as Rop GTPase targets. The Plant Cell, 13, 2841–2856.

87. Yalovsky, S. (2015) Protein lipid modifications and the regulation of ROP GTPase function. J Exp Bot, 66, 1617–1624.

88. Yang, Y., Li, R. and Qi, M. (2000) In vivo analysis of plant promoters and transcription factors by agroinfiltration of tobacco leaves. The Plant Journal, 22, 543–551.

89. Yang, Z. (2002) Small GTPases: versatile signaling switches in plants. Plant Cell, 14 Suppl, S375–388.

90. Zhong, R. and Ye, Z.H. (2003) The SAC domain-containing protein gene family in Arabidopsis. Plant Physiol, 132, 544–555.

91. Zhou, Z., Shi, H., Chen, B., Zhang, R., Huang, S. and Fu, Y. (2015) Arabidopsis RIC1 severs actin filaments at the apex to regulate pollen tube growth. The Plant Cell, 27, 1140–1161.

