## Supplementary material for "Barley resistance and susceptibility to fungal cell entry involve the interplay of ROP signaling with phosphatidylinositol-monophosphates": Suppl. Figures

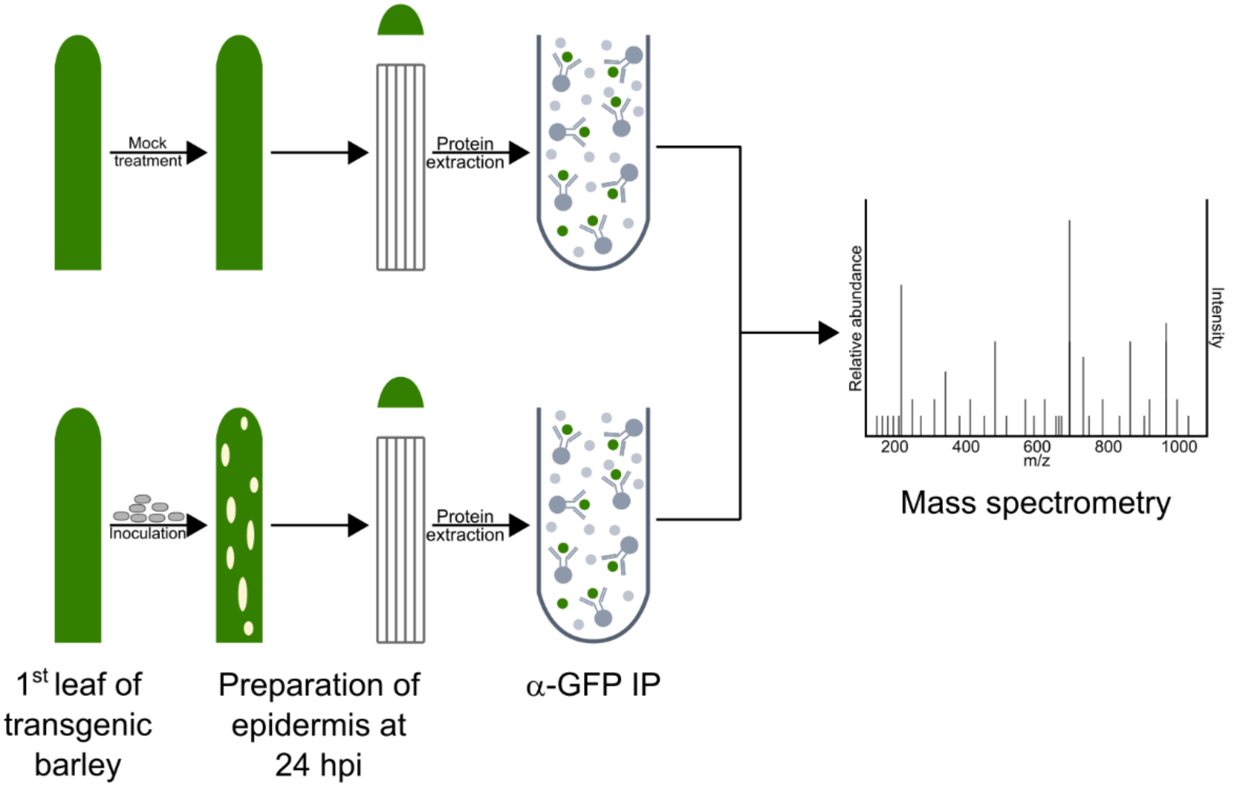


**Supplemental Figure S1: RACB-CA epidermis interactome screening.**
Overview of the *Bh*-infected barley epidermis RACB-CA interactome screening. Transgenic barley plants overexpressing eGFP-RACB-CA, eGFP-RACB-CA-ΔCSIL or free eGFP were either mock-treated or inoculated with *Bh*-spores. At 24 hpi, we produced epidermal peels, extracted total protein, and enriched eGFP-tagged proteins and their putative interactors via α-GFP immunoprecipitation. Pulled-down proteins were identified via LC-MS/MS.

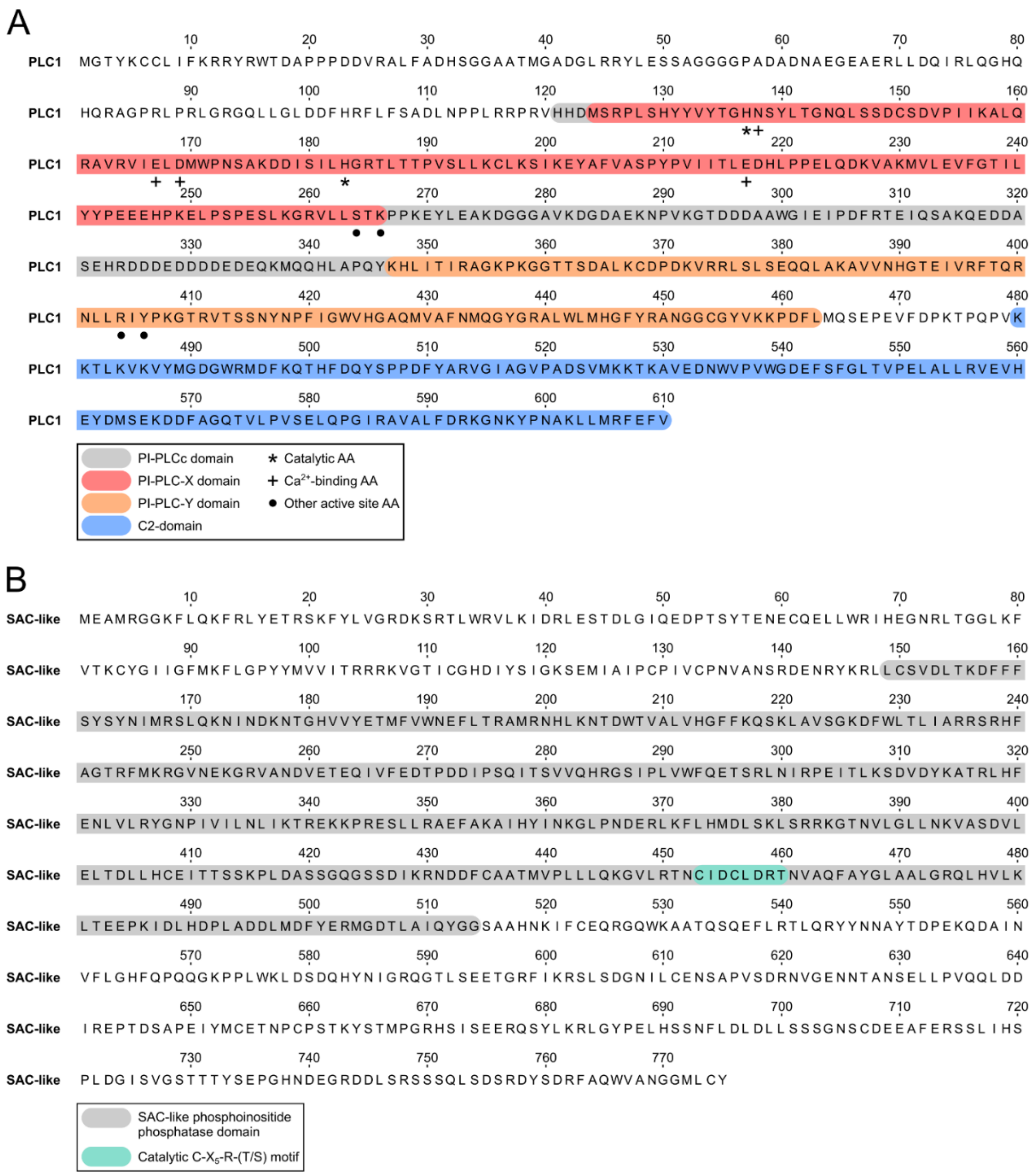


**Supplementary Figure S2: Annotation of functional domains and catalytic amino acids in PLC1 and SAC-like.**

**(a):** Annotation of conserved domains and residues for PLC1, which is based on results from the NCBI conserved domain-search tool (Luo et al. 2020).

**(b):** Annotation of conserved domains and residues for SAC-like, which is based on information from the UniProt database (SAC-like's UniProt identifier: A0A287PS01_HORVV; The UniProt Consortium (2021).

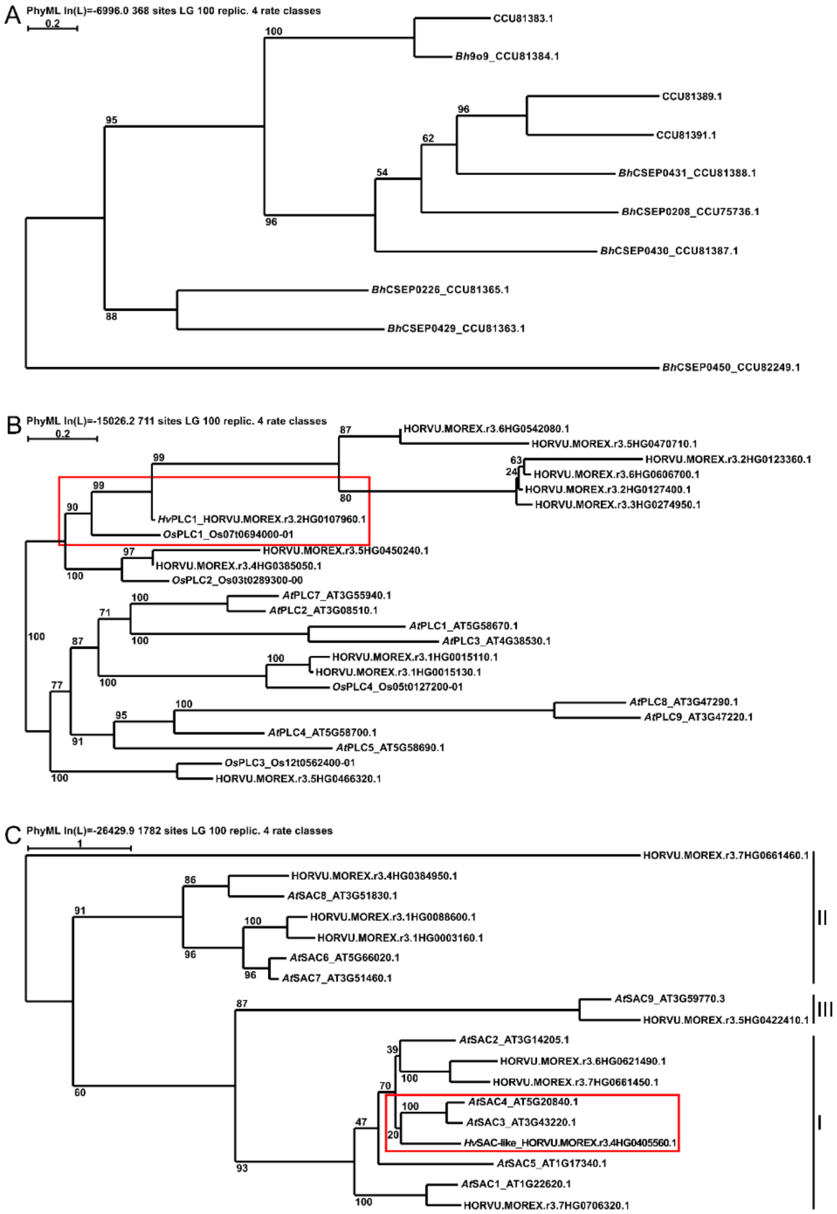


**Supplementary Figure S3: Homologs of 9o9, PLC1 and SAC-like in barley, rice, Arabidopsis and *Bh*.**

Homologous proteins of 9o9 (**a**), PLC1 (**b**) and SAC-like (**c**) were identified by BLASTP searches against the most recent proteomes of *Bh*, barley, rice and Arabidopsis. Maximum-likelihood trees were built in SeaView (V5.0.5, Gouy et al. (2010)) after MUSCLE protein sequence alignment (Edgar, 2004). Values show bootstrap analysis results with 100 iterations. Quality scores are shown above each tree. Branch length indicates the difference between sequences. According to this analysis, red boxes show the closest homologs of PLC1 and SAC-like. The subclades of the Arabidopsis SACs are indicated in Roman numbers in (**c**) on the right according to Zhong and Ye (2003). For more details about these analyses, please see experimental procedures. Please note that for *At*PLC6 no sequence and locus could be identified in the most recent Arabidopsis proteome (Araport11, Cheng et al. (2017).

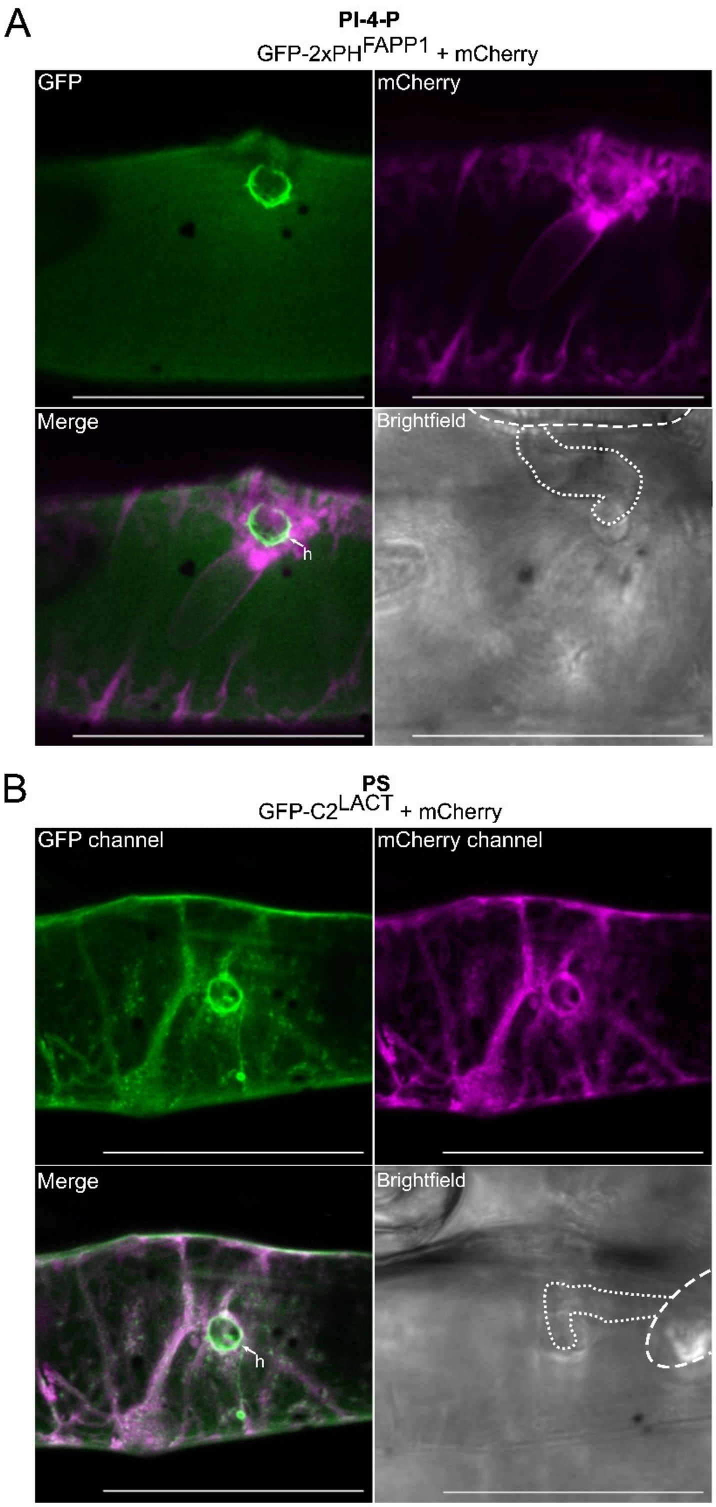


**Supplementary Figure S4: Zoom in magnification of phospholipid marker accumulation at sites of penetration by *B. hordei*.**

**(a,b)** Images of infected cells were taken between 16-20 hpi. High magnification images of non-penetrated *Bh*-colonized (“Haustorium”) cells are shown. Arrows point to haustorial entry points (h). PtdIns4P **(a)** and PtdSer **(b)** markers shown in green were enriched at the haustorial neck region of *Bh*-colonized cells and contrasted by less focal accumulation of cytosolic mCherry shown in magenta. Scale bar: 50 µm.

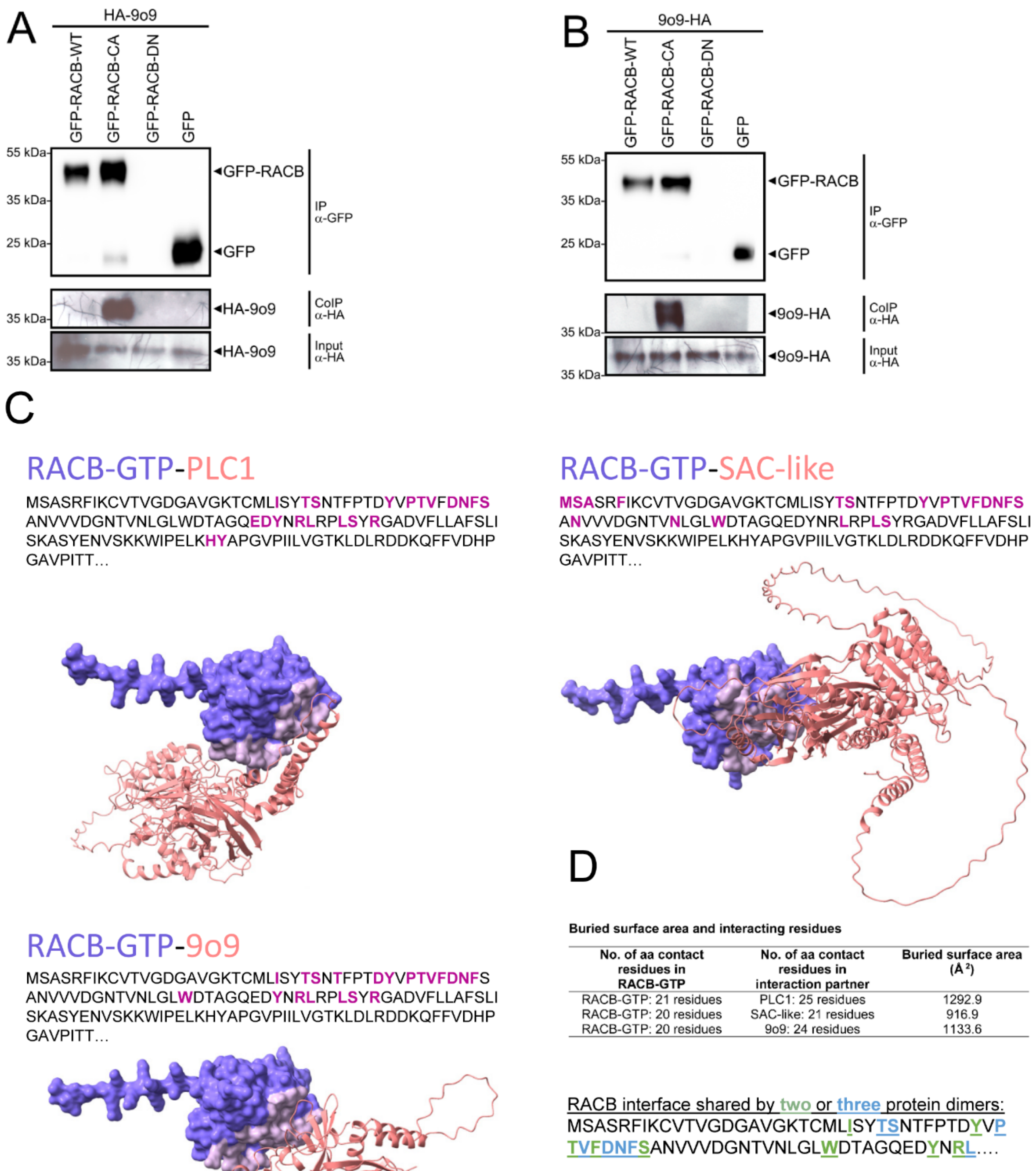


**Supplementary Figure S5: Interaction of 9o9 with RACB *in vivo* and of 9o9, PLC1 and SAC-like with RACB *in silico*.**

(a) and (b) Western blots showing the input and enriched proteins from CoIP experiments after *Agrobacterium tumefaciens*-transformation of *N. benthamiana* plants. GFP-tagged RACB-WT, RACB-CA and free GFP were enriched via αGFP immunoprecipitation. GFP-RACB-DN was not detected in any sample. 9o9 was tagged N-terminally (a) and C-terminally (b)

(c) ALPHAFOLD3 (Abramson et al., 2024) models of protein dimers with the respective amino acid contact sites in RACB-GTP highlighted in magenta in the protein sequence and in grey in the protein model (d) ChimeraX (Pettersen et al., 2021) mediated analysis of contact amino acids and buried intaction interface. 20 or more amino acids on RACB-GTP or the respective binding partner are involved in protein-protein interaction. Protein complexes share the majority of contact amino acids in RACB-GTP (in green or blue in the amino acid sequence), suggesting that protein-protein interaction sites are overlapping.
