## Supplementary material for "Barley resistance and susceptibility to fungal cell entry involve the interplay of ROP signaling with phosphatidylinositol-monophosphates": Suppl. texts

### Supplemental File 1

#### Molecular cloning

For the generation of transgenic barley events, the coding sequence (CDS) of full-length *RACB*-CA (carrying the G15V mutation; Schultheiss et al. (2003)) or *RACB*-CA-ΔCSIL (lacking the last 4 amino acids, but with an artificial stop codon) was amplified by PCR with primers RACB-CA-Esp-Start + RACB-CA-Esp-Stop and RACB-CA-Esp-Start + RACB-CA-ΔCSIL-Esp-Stop, respectively. Both constructs were cloned first into pGGEntL-EP12 and subsequently into pGGInAE-224n_35SP_Ntag GFP_35ST by methods based on GoldenGate cloning techniques (Engler et al., 2008). The eGFP-RACB-CA(-ΔCSIL) fusion constructs and an eGFP-control were re-amplified by PCR using primers GFP-BamHI+2-F + RACB-CA-XmaI-R, GFP-BamHI+2-F + RACB-CA-ΔCSIL-XmaI-R and GFP-BamHI-F + UL1-XmaI-R, respectively, and inserted into pUbifull-AB-M (a derivative of pNOS-AB-M (DNA-Cloning-Service, Hamburg, Germany) with a maize *polyubiquitin* promoter in front of the multiple cloning site) by BamHI + XmaI-mediated restriction ligation. These constructs were transferred by restriction-ligation into p6i-2x35S-TE9 (DNA-Cloning-Service, Hamburg, Germany) using SfiI to create final binary plasmids for stable transformation of barley.

Cloning of the *Nicotiana benthamiana* CoIP constructs for *RACB*-WT, *RACB*-CA and *RACB*-DN was achieved by taking the respective pDONR223-*RACB* plasmids from Weiss et al. (2022) and subcloning the inserts into pGWB6-plasmids (adds N-terminal GFP-tag, later used as free GFP-control; Nakagawa et al. (2007)) via Gateway LR-reactions (Invitrogen, Carlsbad, USA). The CoIP constructs of 9o9 and SAC-like were cloned by PCR-amplifying the respective coding sequences from *Bh*-infected barley epidermal cell cDNA with primers 9o9-attB1-F + 9o9-attB2-R and SAC-attB1-F and SAC-attB2-R for constructs with stop codons and 9o9-attB1-F + 9o9-no-Stop-attB2-R and SAC-attB1-F and SAC-no-STOP-attB2-R for constructs without stop codons, respectively. These amplicons were inserted first into pDONR223 vectors via Gateway BP-reactions and then subcloned into pGWB14 and pGWB15 for HA-tagging (Nakagawa et al., 2007) via Gateway LR-reactions. The CDS of *PLC1* (with and without stop-codon) could not be obtained from cDNA via PCR and was instead synthesized by Twist Bioscience (San Francisco, USA) with 5’ and 3’ Gateway attB-attachment sites and a *Zea mays* codon-optimized nucleotide sequence. Using Gateway BP-reactions, the CDS of *PLC1* (with and without stop codon) were first inserted into pDONR223, and then subcloned into pGWB14 and pGWB15 for HA-tagging (Nakagawa et al., 2007) via Gateway LR-reactions.

The *N. benthamiana* FRET-FLIM constructs for meGFP-RACB-CA, GST-mCherry and CRIB46-mCherry were generated in Trutzenberg et al. (2022). The other constructs from this work were created in the same way. Briefly, fusion constructs comprised of a fluorescent protein, a 10x glycine linker and a protein-of-interest were first linked through Esp3I-mediated GoldenGate cloning (Engler et al., 2008) and subsequently transferred into Gateway-vectors through flanking Gateway attB-sites. For this, the necessary Esp3I-sites and attB-sites were introduced via overhang-PCR, as follows: N-terminal attB1-mCherry-10xGly-Esp3I was amplified with primers B1-mCh-10G-Esp-F + B1-mCh-10G-Esp-R. C-terminal Esp3I-10xGly-mCherry-attB2 was amplified using primers Esp-10G-mCh-B2-F + Esp-10G-mCh-B2-R. N-terminal attB1-9o9-Esp3I, attB1-PLC1-Esp3I and attB1-SAC-like-Esp3I were amplified using primers B1-9o9-Esp-F + B1-9o9-no-Stop-Esp-R, B1-PLC1-Esp-F + B1-PLC1-no-Stop-Esp-R and B1-SAC-Esp-F + B1-SAC-no-Stop-Esp-R, respectively. C-terminal Esp3I-9o9-attB2, Esp3I-PLC1-attB2 and Esp3I-SAC-like-attB2 were amplified using primers Esp-9o9-B2-F + Esp-9o9-B2-R, Esp-PLC1-B2-F + Esp-PLC1-B2-R and Esp-SAC-B2-F + Esp-SAC-B2-R, respectively. After PCR, fusion proteins were assembled through combined Esp3I- and T4 DNA Ligase-mediated Restriction-Ligation cloning (Engler et al., 2008). Here, mCherry-9o9 was fused from attB1-mCherry-10xGly-Esp3I + Esp3I-9o9-attB2, while mCherry-PLC1 was fused from attB1-mCherry-10xGly-Esp3I + Esp3I-PLC1-attB2, and mCherry-SAC-like was fused from attB1-mCherry-10xGly-Esp3I + Esp3I-SAC-like-attB2. In turn, 9o9-mCherry was generated from attB1-9o9-Esp3I + Esp3I-10xGly-mCherry-attB2, while 9o9-PLC1 was generated from attB1-PLC1-Esp3I + Esp3I-10xGly-mCherry-attB2, and SAC-like-mCherry was generated from attB1-SAC-like-Esp3I + Esp3I-10xGly-mCherry-attB2. After assembly, all fusion constructs were first inserted into pDONR223 via Gateway BP-cloning and subsequently transferred into pGWB2 (Nakagawa et al., 2007) via Gateway LR-reactions.

The barley overexpression construct of 9o9 was cloned by amplifying the coding sequence of *9o9* via PCR from *Bh*-infected barley epidermal cell cDNA with primers 9o9-F + 9o9-R. This PCR product was phosphorylated using a polynucleotide kinase reaction (Thermo Fisher Scientific, Waltham, USA) and inserted into an empty SmaI-digested pGY1 backbone using a blunt-end T4 DNA Ligation (Thermo Fisher Scientific, Waltham, USA). The overexpression constructs for PLC1 and SAC-like were cloned by performing LR-reactions (Invitrogen, Carlsbad, USA) with a Gateway-compatible pGY1 vector (generated in Engelhardt et al. (2022)) and the pDONR223-PLC1 and pDONR223-SAC-like entry plasmids from above.

The RNAi-silencing constructs for 9o9, PLC1 and SAC-like were generated by first amplifying parts of their coding sequences with primers 9o9-RNAi-XbaI-F + 9o9-RNAi-SalI-R, PLC1-RNAi-XbaI-F + PLC1-RNAi-SalI-R and SAC-RNAi-XbaI-F + SAC-RNAi-SalI-R. Fitting nucleotide sequence stretches with a high probability for gene-specific RNAi were predicted using the si-Fi software (Lück et al., 2019). PCR amplicons were inserted into pIPKTA38 vectors (Douchkov et al., 2005) in 3'-to-5' orientation using SalI + XbaI-mediated classical cloning (Thermo Fisher Scientific, Waltham, USA). From pIPKTA38, the sequences were transferred into the destination vector pIPKTA30N (Douchkov et al., 2005) using Gateway LR-reactions, creating antisense-hairpin-sense RNAi-constructs.

For the yeast secretion assay, full-length 9o9, a putative signal peptide-deficient mutant (9o9-Δ15, lacking amino acids 2-15) and the signal peptide of the *Arabidopsis* RLK LORE ($SP^{LORE}$) were amplified via PCR with primers 9o9-EcoRI-F + 9o9-no-Stop-NotI-R, 9o9-d15-EcoRI-F + 9o9-no-Stop-NotI-R or SP (Rapazote-Flores *et al.*)-EcoRI-F + SP (Rapazote-Flores *et al.*)-NotI-R. Following an EcoRI + NotI digestion of the inserts and the pSmash plasmid, which contained a signal peptide-lacking mutant of the yeast invertase SUC2 (Goo et al., 1999), constructs were assembled via sticky-end T4 DNA ligation (Thermo Scientific, Waltham, USA). The $SP^{9o9}$-SUC2 construct (amino acids 1-20) was generated by PCR- and Esp3I-mediated partial deletion of 9o9 from pSmash-9o9. For this, primers SP(9o9)-deletion-Esp3I-F and SP(9o9)-deletion-Esp3I-R were used to amplify the full pSmash backbone and signal peptide of 9o9, but missing the rest of the 9o9 sequence. The backbone was fused using a combined Esp3I- and T4 DNA Ligase-mediated Restriction-Ligation reaction (Engler et al., 2008).

For recombinant protein expression in *E. coli*, RACB-WT was amplified via PCR from pDONR223-RACB-WT (from Weiss et al. (2022)) with primers RACB-SalI-F + RACB-NotI-R and ligated through SalI + NotI-mediated classical cloning into pGEX-6P-1 (GE-Healthcare, Chicago, USA). The RACB-5Q mutant was generated by amplifying the backbone of the finished pGEX-RACB-plasmid with primers RACB-5Q-F + RACB-5Q-R, which exchanged the 5 lysines in RACB's PBR to 5 glutamines. Since the amplicon was still linearized, it was phosphorylated using a polynucleotide kinase reaction and circularized using a blunt-end T4 DNA ligation (both Thermo Fisher Scientific, Waltham, USA). 9o9, PLC1 and SAC-like were first PCR amplified with primers 9o9-NotI-F + 9o9-SalI-R, PLC1-NotI-F + PLC1-SalI-R and SAC-NotI-F + SAC-SalI-R using their pDONR223-plasmids from above as templates. Subsequently, they were inserted into pMAL-c5X (NEB, Ipswich, USA) by SalI + NotI-mediated classical cloning.

Cloning of RACB-CA-5Q and meGFP-RACB-CA-5Q for transient barley transformation experiments was done by taking the pGY1-RACB-CA and pGY1-meGFP-RACB-CA plasmids from Schultheiss *et al.*, 2003, and Engelhardt *et al.*, 2022, respectively, and repeating the cloning procedure as described for pGEX-RACB-5Q above. pGY1-mCherry-RIC171 and empty pGY1-mCherry were taken from Trutzenberg *et al.*, 2022. Empty pGY1, pGY1-CFP and pGY1-meGFP plasmids were taken from Schultheiss *et al.*, 2003, McCollum *et al.*, 2020, and Trutzenberg *et al.*, 2022, respectively.

Cloning of RACB-5Q yeast expression plasmids was performed by taking the pGBKT7-RACB-WT/CA/DN-deltaC plasmids from Trutzenberg *et al.*, 2022, and repeating the cloning procedure as described for pGEX-RACB-5Q above. Empty pGADT7- and pGBKT7-vectors as well as pGADT7-RIC171 were also taken from Trutzenberg *et al.*, 2022.

Cloning of the barley overexpression constructs of the anionic phospholipid markers was achieved by a mixture of GoldenGate and Gateway cloning techniques. To create pGY1-GFP-2xPH$^{FAPP1}$, a GoldenGate-compatible pGY1-Esp3I amplicon was generated by taking empty pGY1 as template (Schweizer et al., 1999) and amplifying it using primers pGY1-Esp-F + pGY1-Esp-R to attach Esp3I-sites. GFP was amplified from pGY1-meGFP (Engelhardt et al., 2022) with primers FP-Esp-F + FP-Esp-R. The two copies of PH$^{FAPP1}$ were amplified from P21Y (Simon et al. (2014); clone was ordered from NASC (ID N2106328)) with primers FAPP1-Esp-1st-F + FAPP1-Esp-1st-R and FAPP1-Esp-2nd-F + FAPP1-Esp-2nd-R, respectively. The four amplicons were combined in a single GoldenGate restriction-ligation reaction using Esp3I and T4 DNA ligase (both Thermo Fisher Scientific, Waltham, USA; Engler et al. (2008)) to create the final overexpression construct. pGY1-GFP-C2$^{LACT}$ was cloned similarly, amplifying C2$^{LACT}$ from pB7m34GW-pUBQ10::Citrine-C2$^{LACT}$ (Platre et al. (2018); a gift from Yvon Jaillais (ENS Lyon) with primers LACT-Esp-F + LACT-Esp-R and combining it with pGY1-Esp and GFP-Esp from above in a GoldenGate restriction-ligation (Engler et al., 2008). Cloning of pGY1-mCitrine-2xML1N was achieved by amplifying the full mCitrine-2xML1N cassette with attB-sites from pGWB501-UBQ10::mCitrine-2xML1N (Hirano et al. (2017b); a gift from Masa Sato (Kyoto Prefectural University)) with primers ML1N-attB-F + ML1N-attB-R. This construct was first introduced into pDONR223 via a Gateway BP-reaction and then transferred into Gateway-compatible pGY1 (Engelhardt et al., 2022) via a Gateway LR-reaction to create the barley overexpression construct. pGY1-GFP-2xPH$^{PLC}$ was cloned by amplifying 2xPH$^{PLC}$ from P24Y (Simon et al. (2014); clone ordered from NASC (ID N2106329)) with primers PHPLC-attB-F + PHPLC-attB-R and inserting it into pDONR223 via a Gateway BP-reaction. From there, 2xPH$^{PLC}$ was transferred into Gateway-compatible pGY1-meGFP (Engelhardt et al., 2022) using a Gateway LR-reaction to create the barley localization construct.

After every significant cloning step, insert and fusion construct sequence identities were confirmed by Sanger sequencing (Eurofins Genomics, Ebersberg, Germany). All primers can be found in Supplemental Table 2. All gene identifiers can be found in Supplemental Table 3.

**Generation and selection of transgenic barley**

The transgenic barley lines from this study were largely produced in a way as those described in (Weiss et al., 2022). Final binary plasmids containing the constructs-of-interest (see section molecular cloning) were used to transform *Agrobacterium tumefaciens* strain AGL1 (Lazo et al., 1991). Transgenic barley plants were generated by *A. tumefaciens*-mediated DNA-transfer to immature barley embryos (cultivar Golden Promise) following the protocol from Hensel et al. (2009). Transgenic plants were identified by HYGROMYCIN PHOSPHOTRANSFERASE-specific PCR, which yielded 27 events for eGFP-RACB-CA (BG654 E1-E29; E16 and E29 were not transgenic), 14 events for eGFP-RACB-CA-ΔCSIL (BG655 E1-E15; E13 was not transgenic) and 17 events for eGFP (BG656 E1-E17). All events were propagated under normal growth conditions in the greenhouse. Since all offspring were segregating in generation $T_1$, individual plants were screened for transgene integration. This was done via selection using Hygromycin B (as described in Weiss et al. (2022)), GFP-fluorescence microscopy and routine α-GFP Western blotting. After screening, ten transgene-expressing plants of two events per construct were picked for an additional round of propagation: we chose BG654 E02 and E12 for eGFP-RACB-CA, BG655 E01 and E10 for eGFP-RACB-CA-ΔCSIL and BG656 E01 and E06 for eGFP. Since the descendants of these events remained segregating in generation $T_2$ and no homozygous plants could be obtained, all offspring were screened for transgene expression as described above before being used in any experiment. All primers can be found in Supplemental Table 2. All antibodies can be found in Supplemental Table 4.

**Transient transformation of *Nicotiana benthamiana* plants**

Six to seven weeks old *Nicotiana benthamiana* plants were transformed via *Agrobacterium tumefaciens* (strain GV3101 pMP90; Lazo et al. (1991)) with a protocol adapted from Yang et al. (2000). For that, chemically-competent *Agrobacterium tumefaciens* were first transformed with binary pGWB plasmids containing constructs-of-interest (see molecular cloning). Agrobacteria containing the respective pGWB plasmids were then struck on LB-agar plates containing 10 µg/ml rifampicin, 30 µg/ml gentamicin and 50 µg/ml kanamycin and grown for 3 d at 28 °C. Afterwards, 2.5 ml of induction medium (1 g/l $NH_4Cl$, 0.3 g/l $MgSO_4$-7*$H_2O$, 0.15 g/l KCl, 0.01 g/l $CaCl_2$, 0.0025 g/l $FeSO_4$-7*$H_2O$, 3 g/l $K_2HPO_4$, 1 g/l $NaH_2PO_4$, 10 g/l D-glucose, 20 mM MES pH 5.5 (2-(N-morpholino)ethanesulfonic acid) supplied with 100 µM acetosyringone, 30 µg/ml gentamicin and 50 µg/ml kanamycin were inoculated with growing agrobacteria clones and incubated overnight in a 28 °C shaker. Next day, agrobacteria were harvested using centrifugation at 3000 g for 2 min, then washed twice with 1 ml infiltration medium (10 mM $MgSO_4$, 10 mM MES pH5.5) before being taken up in 1 ml infiltration medium with 150 µM acetosyringone. Bacterial density in the samples was measured and adjusted to an $OD_{600}$ of 0.5 using infiltration medium with acetosyringone. For protein co-expression *in planta*, agrobacteria containing the corresponding plasmids were mixed with agrobacteria containing a plasmid coding for the silencing suppressor p19 (Voinnet et al., 2003) in a 1:1:1 ratio. Following a 1 h incubation at room temperature, bacterial solutions were infiltrated into the abaxial side of fully expanded *Nicotiana benthamiana* leaves using a syringe without a needle. Infiltrated plants were placed in climate chambers operating under long-day conditions (see above). CoIP and FRET-FLIM experiments were performed 2 d after infiltration.

**Transient transformation of barley**

Epidermal cells of wildtype barley *cv.* Golden Promise were transiently transformed via particle bombardment with a protocol adapted from (Schweizer et al., 1999). For one reaction, 11 µl of a spherical gold nanoparticle solution (27.5 µg/ml; 1 µm diameter; Bio-Rad, Hercules, USA) were mixed with plasmids encoding constructs-of-interest (see molecular cloning). For transformation markers, such as free fluorophores or GUS+ (Vickers et al., 2003), 0.5 µg plasmid was used per replicate, while for all other constructs 1 µg/replicate of plasmid was taken. The combined volume of gold and plasmids was doubled with an equal amount of 2.5 M $CaCl_2$, followed by an addition of 3.33 µl of a 20 mg/ml Protamine solution. This solution was incubated for 30 minutes at room temperature with mixing every 10 minutes. Afterwards, the gold particles were pelleted by centrifugation at max. speed for 10 s and the supernatant was removed. The gold particles were washed twice with 400 µl 100% ethanol, resuspended in 6 µl of 100% ethanol and being spotted on Macrocarriers (Bio-Rad, Hercules, USA) for particle bombardment. For a single transformation reaction, three primary barley leaves were detached and placed on 0.8% water-agar plates with the adaxial side facing up. These leaves were transformed with a PDS-1000/HE™ (Bio-Rad, Hercules, USA) particle gun, operating with 900 psi rupture discs and 26 inHg. To analyse *Bh* susceptibility, at least five biological replicates were created; all other experiments used at least two replicates.

**Analysis of *Bh* penetration efficiency**

The *Bh* susceptibility assessment of transgenic plants was conducted in events BG654 E2 for eGFP-RACB-CA, BG655 E1 for eGFP-RACB-CA-ΔCSIL and BG656 E1 for eGFP. All plants were selected, transgene-expressing offspring in generation $T_1$. For this experiment, three primary leaves per construct were cut and placed on 0.8% water-agar plates. All plates were inoculated with 10-20 *Bh* spores/mm$^2$ and incubated under normal growth conditions for 40 h. Afterwards, the leaves were fixed and destained in 70% (v/v) ethanol before analysis. For the *Bh* susceptibility assay with transiently transformed plants, leaves were first transformed with either plasmid for gene overexpression (pGY1, Schweizer et al. (1999) or RNAi-mediated silencing (pIPKTA30N, Douchkov et al. (2005)) as described above. All constructs (see molecular cloning) were co-transformed with the transformation marker GUS+ (a gift from Claudia Vickers, Addgene plasmid #64402, Vickers et al. (2003)) to later identify transformed cells. One day (overexpression) or two days (RNAi) after transformation, the bombarded leaves were inoculated with 100-130 *Bh* spores/mm$^2$ and kept under normal growth conditions (see above). At 40 hpi, the transformed leaves were placed in plates containing a GUS-staining solution (0.1 M $Na_2HPO_4/NaH_2PO_4$ buffer pH 7, 0.01 M sodium EDTA, 0.005 M potassium hexacyanoferrat (II), 0.005 M potassium hexacyanoferrate  (Mackey *et al.*), 0.1% (v/v) Triton X-100, 20% (v/v) methanol, 0.5 mg/ml X-gluc (1,5-bromo-4-chloro-3-indoxyl-β-D-glucuronic acid, cyclohexyl ammonium salt (Carbosynth, Bratislava, Slovak Republic); Schweizer et al. (1999)). Following a 3 min vacuum infiltration, the leaves were incubated for at least 4 h at 37 °C in the dark until sufficient GUS-staining was achieved. Samples were destained in 70% (v/v) ethanol. A combination of fluorescence and transmission light microscopy analyzed destained leaves. First, fungal structures were stained with an ink solution (10% (v/v) Pelikan Ink 4001 (Pelikan, Hannover, Germany), 25% (v/v) acetic acid; transgenic plants only) or with a Calcofluor White staining solution (0.3% (w/v) Calcofluor White M2R (F-3543, Sigma-Aldrich, St. Louis, USA), 50 mM TRIS, 2% (v/v) Tween20, pH 9; transiently transformed leaves only) for better visibility. On transiently transformed leaves, only GUS-stained epidermal A and B cells were considered for analysis, while on stable transgenic leaves, all epidermal A and B cells were considered. In the respective cells, haustorium establishment was evaluated as successful penetration, whereas stopped fungal growth after appressorium formation was counted as successful plant defense. About 100 plant-fungus interaction sites were analyzed per leaf. For each leaf, the *Bh* penetration efficiency was calculated by dividing the number of haustoria by the sum of all evaluated interaction sites on this leaf. For Fig. 1 C, these values were divided by the average penetration efficiency of the eGFP control. In the transient assay, the *Bh* penetration efficiency was determined separately for each plasmid combination by dividing the number of all established haustoria by the sum of all evaluated plant-fungus interactions of one combination. At least 5 independent biological replicates were performed per experiment, with a minimum of 50 evaluated plant-fungus interactions per plasmid combination per replicate. For Fig. 4 A-D, all *Bh* penetration efficiencies were divided by the average penetration efficiency of their respective controls. All data was plotted in GraphPad Prism V8.0 (GraphPad Software, San Diego, USA) and analyzed by a One-way ANOVA with Tukey's HSD (transgenic plants) or Student's *t*-tests (transiently transformed plants) against an $\alpha$ of 0.05, which were calculated in RStudio V1.2.5033.

**FRET-FLIM measurements**

FRET-FLIM measurements were performed with an FCS/FLIM-FRET/rapidFLIM upgrade kit (Picoquant, Berlin, Germany) used in tandem with an Olympus (Tokyo, Japan) FV3000 mounted on an IX63 stand. This method was adapted from (Weidtkamp-Peters and Stahl, 2017). In *Nicotiana benthamiana*, FRET-FLIM measurements were carried out at the cell periphery of two adjacent epidermal cells, which both showed co-expression of meGFP- and mCherry-fusion proteins. In barley, FRET-FLIM measurements were performed in transiently transformed barley epidermal cells as described in Trutzenberg *et al.*, 2022. All cells were observed with an UPLSAPO60XW 60x/NA 1.2/WD 0.28 water immersion objective (Olympus, Tokyo, Japan) and 4x zoom to reach the Nyquist-criterion. GFP-lifetime measurements were conducted using the time-correlated single-photon counting (TCSPC) method from the Picoquant kit. For this, meGFP was excited with a pulsed 485 nm diode laser (LDH-D-C-485, pulse rate: 40 MHz) and its emission was detected with two photon-counting PMA Hybrid 40 detectors operating two TCSPC modules (TimeHarp 260 PICO Dual, TimeHarp 260 NANO Dual). Image resolution was 512*512 pixels, while TCSPC resolution was 25 ps. At least 500 photons were collected in the brightest pixel for one measurement. Before analysis, regions of interest (ROIs) were used to select image areas in focus and showing fluorescence. Lifetime-decay-fitting and analysis was done with the SymPhoTime 64 software V2.6 (PicoQuant, Berlin, Germany) using a software-calculated instrument response factor and an n-exponential reconvolution with n = 2. Only measurements with $\chi^2$-values between 1.0 and 2.0 and positive amplitudes were considered for quality control. We measured the intensity-weighted average GFP-lifetime τ and plotted this in GraphPad Prism V8.0 (GraphPad Software, San Diego, USA). Statistical analysis was done in RStudio V1.2.5033 using Wilcoxon Rank-Sum tests against an α of 0.05 with Bonferroni-correction.

**Yeast secretion assay**

The yeast secretion assay was adapted from Krijger et al. (2008). Yeast strain Y02321 (BY4741; MATa; *his3*Δ1; *leu2*Δ0; *met15*Δ0; *ura3*Δ0; YIL162w::kanMX4; from EUROSCARF, Oberursel, Germany) was turned chemically competent with a protocol adapted from Gietz and Woods (2002). In brief, yeast cells were struck on YPD plates (1% (w/v) yeast extract, 2% (w/v) peptone, 2% (w/v) D-glucose, 2% (w/v) agar) and incubated for 3 d at 30 °C. After that, one newly grown colony was used to inoculate 5 ml liquid YPD medium and the culture was incubated overnight in a 30 °C shaker. The overnight culture was then used to inoculate 100 ml liquid YPD medium, and this culture was grown for 5 h at 30 °C in a shaker. Subsequently, the culture was divided into two 50 ml fractions, centrifuged for 5 min at 700 g to pellet the cells and remove the medium. After two washing steps with 30 ml sterile $H_2O$ and centrifugation as above, each pellet was resuspended in 1.5 µl TELiAc buffer (11 mM Tris-HCl pH 7.5, 1.1 mM EDTA, 110 mM lithium acetate) and the solutions were transferred into 2 ml tubes. The cells were again pelleted via centrifugation at 18000 g for 15 s, after which the supernatants were removed, 600 µl TELiAc buffer were added and the solutions were combined in one tube after resuspension.

For transformation, 100 ng of pSmash-plasmids (see molecular cloning) were mixed with 10 µl of boiled salmon sperm carrier DNA (Invitrogen, Carlsbad, USA) and 50 µl of chemically competent cells. Followingly, 500 µl of sterile PEG/LiAc buffer (40% PEG3350, 10 mM Tris-HCl pH 7.5, 1 mM EDTA, 100 mM lithium acetate) were added and the solutions were mixed by pipetting. After a 30 min incubation at 30 °C with careful mixing every 10 min, 20 µl DMSO were added and the solutions were mixed by pipetting. Following a 15 min incubation at 42 °C with constant shaking, the cells were centrifuged at 18000 g for 15 s and the supernatants were removed. The cells were washed once with 1 ml of TE buffer (10 mM Tris-HCl pH 7.5, 1 mM EDTA) and centrifugation as above, after which the supernatants were removed and the cells were resuspended in 50 µl TE buffer. To select for transformed cells, all samples were plated on SD/-L plates containing 2% D-glucose (6.7 g/l yeast nitrogen base without amino acids, 670 mg/l Complete Supplement Mixture without leucine, 2% (w/v) agar, 2% (w/v) D-glucose; all from Formedium, Hunstanton, UK) and grown for 3 d at 30 °C. To test secretion, three growing colonies per sample were first resuspended in 100 µl sterile $H_2O$ and then diluted from 1 (undiluted) to 1/3125 (equals six 1/5 dilution steps). From each dilution step, 7.5 µl were dropped on both a single SD/-L + 2% D-glucose plate and a single SD/-L + 2% raffinose + 2 µg/ml Antimycin A (Sigma-Aldrich, St. Louis, USA) plate to compare secretion between samples. All plates were incubated for at least 3 d at 30 °C until yeast growth could be observed. In this experiment, yeast growth on SD/-L + 2% raffinose + 2 µg/ml Antimycin A plates confirmed secretion of the construct, since a sucrose-hydrolyzing SUC2 enzyme lacking its N-terminal signal peptide was fused in frame to the C-terminus of a gene-of-interest. If the gene-of-interest was secreted, SUC2 would thus also be targeted to the extracellular medium, where it hydrolyzes sucrose. Yeast strain Y02321 is a sucrose-auxotroph and can only grow, when extracellular sucrose is hydrolyzed to D-glucose and D-fructose. Here, raffinose (a tri-saccharide of sucrose + D-galactose) instead of sucrose was used to increase selection pressure. Antimycin A (an inhibitor of oxidative respiration) was included to force yeast cells to rely on fermentation (Krijger et al., 2008). All primers can be found in Supplemental Table 2.

**RT-qPCR**

Gene expression of *9o9*, *PLC1* and *SAC-like* was measured using RT-qPCR. For each biological replicate, 6 pots of 20 wildtype barley (*cv.* Golden Promise) seeds were grown in a climate chamber (for conditions, see above). After growing for 7 days, 3 pots were inoculated with 100-130 *Bh* spores/mm$^2$ and placed in a different climate chamber to avoid contamination. At the respective timepoints (6, 12 and 24 hpi), the abaxial epidermis was collected from all plants of one pot and stored in a 2 ml tube in liquid $N_2$. Each tube contained two 4 mm glass beads for later sample homogenization. Uninfected control samples were collected in the same fashion at the indicated timepoints. All plant samples were stored at -80 °C until homogenization. *Bh* spores were collected by placing infected leaves (age roughly 7 dpi, from *Bh* propagation) in a 15 ml tube and shaking until all spores had fallen off. Spores from 20 *Bh*-infected leaves were collected in this fashion for one biological replicate. Spores sticking to the side of the tube were washed down with 1 ml aqueous 0.05% (V/V) Tween20 solution and inversion of the tube. The spore solution was transferred to a 2 ml tube and the liquid was removed by centrifugation at 6000 g for 10 s. Two 4 mm glass beads and 100 mg 0.1 mm glass beads were added, the sample was frozen in liquid $N_2$ and stored at -80 °C until homogenization. Sample homogenization was carried out using a TissueLyser II (QIAGEN, Hilden, Germany), which ran twice for 1 min at 30 Hz with a cooling step in liquid $N_2$ in between. For RNA extraction, 1 ml of TRIzol (Thermo Fisher Scientific, Waltham, USA) was added to each sample, followed by thorough vortexing. Afterwards, the Direct-zol RNA Mini Prep Kit (R2052, Zymo Research Europe, Freiburg, Germany) including the on-column DNAse treatment was used and the manufacturer's were followed. The extracted RNA was directly used for cDNA synthesis, which was carried out with the RevertAid RT Kit (Thermo Fisher Scientific, Waltham, USA), using 500 ng of RNA as template and the optional boiling step for GC-rich samples as recommended by the manufacturer. cDNA was stored at -20 °C until use. For RT-qPCR, the AriaMx Real-time PCR System (Agilent, Santa Clara, USA) and the Takyon™ Low ROX SYBR 2x MasterMix dTTP Blue Kit (Kaneka Eurogentec S.A., Seraing, Belgium) were used. Threehundred nM forward and reverse primers and 10 ng cDNA were used for one reaction. Each RT-qPCR reaction was prepared and run as recommended by the manufacturer (Kaneka Eurogentec S.A., Seraing, Belgium). To avoid inter-run variances, all samples and timepoints were run in triplicate on the same 96-well plate for each gene. After identification, outliers were excluded from analysis, according to differences in melt-curves and discrepancies between technical replicates. For each valid measurement, the cycle threshold (Ct) values were exported as ΔRn values, which included automatic software-based correction steps from the Agilent AriaMx software V1.8 (Agilent, Santa Clara, USA). Prior to gene expression measurements, primer tests were conducted for each primer pair. For this, samples of all timepoints and replicates were pooled according to treatments. All *Bh*-containing samples (lacking the fully untreated plant control) and all plant samples (lacking the *Bh* spore sample) were pooled separately. Each pool was diluted in 1/10 dilution steps from initial 10 ng/µl down to 0.01 ng/µl. All barley primer pairs were tested on the plant samples, while *Bh* primers were tested on the *Bh* pool. Primer efficiencies were calculated by the Agilent AriaMx V1.8 software (Agilent, Santa Clara, USA). Primer specificity was confirmed using Sanger sequencing of RT-qPCR amplicons (Genewiz Europe, Leipzig, Germany). Gene expression data was calculated as normalized fold changes in Microsoft Excel 2016 (Microsoft, Redmond, USA) according to Pfaffl (2001). Experimentally determined primer efficiencies had to be used, as all primer pairs except the one for *β-Tub2* showed an efficiency deviating from 100% amplification rate. As housekeeping genes, *ubiquitin-conjugating enzyme 2* (*UBC2*) was used for barley, while *β-tubulin 2* (*β-TUB2*) was used for *Bh* (Sherwood and Somerville, 1990; Rapacz et al., 2012; Schnepf et al., 2018). Data was plotted and analysed in GraphPad Prism V8.0 (GraphPad Software, San Diego, USA). Statistical analysis compared normalized gene expression in infected leaves to that of corresponding uninfected leaves via *t*-tests against an α of 0.05 with Holm-Sidak correction for multiple testing on $\log_{10}$-transformed data. All primers can be found in Supplemental Table 2. All genes can be found in Supplemental Table 3.

**Bioinformatic analyses**

The protein domains of 9o9, PLC1 and SAC-like were predicted by comparing the respective amino acid sequences against the UniProt database (The UniProt Consortium 2021) and by feeding the sequences into the NCBI CD-search algorithm (https://www.ncbi.nlm.nih.gov/Structure/cdd/wrpsb.cgi; Lu et al. (2020)). Identified catalytic amino acid and conserved domains were highlighted in the protein sequences of PLC1 and SAC-like via Inkscape V1.2 (https://inkscape.org/).

Signal peptide prediction for 9o9 was performed with SignalP V6.0 (Teufel et al., 2022).

Homologs of 9o9, PLC1 and SAC-like in *Blumeria hordei*, barley, *Arabidopsis thaliana* and rice were identified by blasting the respective protein sequences against the following proteomes: *Blumeria hordei*: isolate DH14 (Frantzeskakis et al., 2018); barley: cultivar Morex V3 (Mascher, 2021); *Arabidopsis thaliana*: Araport11 (Cheng et al., 2017); rice: IRGSP 1.0 (Kawahara et al., 2013). Blasting 9o9 revealed and unusually high number of potentially homologous proteins, which is why candidates were filtered according to the following criteria: putative homologs had to be identified with an error value smaller than 0.05, an aligned sequence coverage higher than 30% and a sequence identity above 30%.

For phylogenetic analysis and PhyML tree-building, all identified homologs of 9o9, PLC1 and SAC-like were first aligned with the MUSCLE-alignment tool running with default parameters in SeaView V5.0.5 (Edgar, 2004; Gouy et al., 2010). From the multiple-sequence alignments, PhyML trees were built in SeaView with the following parameters: LG-model, bootstrap analysis with 100 replicates, model-given amino-acid equilibrium frequencies, no invariable sites, optimized across site rate variation, nearest-neighbor interchange for tree searching and five random starts. Calculated trees were exported and modelled in Inkscape V1.2 (https://inkscape.org/).

The barley reference transcript database (BaRTD (Mascher et al., 2017; Rapazote-Flores et al., 2019)) was used to identify *PLC1* and *SAC-like* expression patterns in different barley tissues. Matching transcripts were identified by blasting the nucleotide sequences of *PLC1* and *SAC-like* against the database. The gene expression levels were exported from the website and plotted in GraphPad Prism V8.0 (GraphPad Software, San Diego, USA).

A list of all genes and their identifiers can be found in Supplemental Table 3.

#### References for supplementary figures, tables and methods


**Abramson, J., Adler, J., Dunger, J., Evans, R., Green, T., Pritzel, A., Ronneberger, O., Willmore, L., Ballard, A.J., Bambrick, J., Bodenstein, S.W., Evans, D.A., Hung, C.-C., O'Neill, M., Reiman, D., Tunyasuvunakool, K., Wu, Z., Žemgulytė, A., Arvaniti, E., Beattie, C., Bertolli, O., Bridgland, A., Cherepanov, A., Congreve, M., Cowen-Rivers, A.I., Cowie, A., Figurnov, M., Fuchs, F.B., Gladman, H., Jain, R., Khan, Y.A., Low, C.M.R., Perlin, K., Potapenko, A., Savy, P., Singh, S., Stecula, A., Thillaisundaram, A., Tong, C., Yakneen, S., Zhong, E.D., Zielinski, M., Žídek, A., Bapst, V., Kohli, P., Jaderberg, M., Hassabis, D. and Jumper, J.M.** (2024) Accurate structure prediction of biomolecular interactions with AlphaFold 3. *Nature*, **630**, 493-500.

**Cheng, C.Y., Krishnakumar, V., Chan, A.P., Thibaud-Nissen, F., Schobel, S. and Town, C.D.** (2017) Araport11: a complete reannotation of the Arabidopsis thaliana reference genome. *Plant J*, **89**, 789-804.

**Douchkov, D., Nowara, D., Zierold, U. and Schweizer, P.** (2005) A high-throughput gene-silencing system for the functional assessment of defense-related genes in barley epidermal cells. *Molecular Plant-Microbe Interactions*, **18**, 755-761.

**Engler, C., Kandzia, R. and Marillonnet, S.** (2008) A one pot, one step, precision cloning method with high throughput capability. *PLoS One*, **3**, e3647.

**Essen, L.-O., Perisic, O., Cheung, R., Katan, M. and Williams, R.L.** (1996) Crystal structure of a mammalian phosphoinositide-specific phospholipase Cδ. *Nature*, **380**, 595-602.

**Essen, L.-O., Perisic, O., Katan, M., Wu, Y., Roberts, M.F. and Williams, R.L.** (1997) Structural mapping of the catalytic mechanism for a mammalian phosphoinositide-specific phospholipase C. *Biochemistry*, **36**, 1704-1718.

**Goo, J.H., Park, A.R., Park, W.J. and Park, O.K.** (1999) Selection of Arabidopsis genes encoding secreted and plasma membrane proteins. *Plant molecular biology*, **41**, 415-423.

**Gouy, M., Guindon, S. and Gascuel, O.** (2010) SeaView version 4: a multiplatform graphical user interface for sequence alignment and phylogenetic tree building. *Molecular biology and evolution*, **27**, 221-224.

**Hensel, G., Kastner, C., Oleszczuk, S., Riechen, J. and Kumlehn, J.** (2009) Agrobacterium-mediated gene transfer to cereal crop plants: current protocols for barley, wheat, triticale, and maize. *Int J Plant Genomics*, **2009**, 835608.

**Julkowska, M.M., Rankenberg, J.M. and Testerink, C.** (2013) Liposome-Binding Assays to Assess Specificity and Affinity of Phospholipid–Protein Interactions. In *Plant Lipid Signaling Protocols* (Munnik, T. and Heilmann, I. eds). Totowa, NJ: Humana Press, pp. 261-271.

**Kawahara, Y., de la Bastide, M., Hamilton, J.P., Kanamori, H., McCombie, W.R., Ouyang, S., Schwartz, D.C., Tanaka, T., Wu, J. and Zhou, S.** (2013) Improvement of the Oryza sativa Nipponbare reference genome using next generation sequence and optical map data. *Rice*, **6**, 1-10.

**Krijger, J.-J., Horbach, R., Behr, M., Schweizer, P., Deising, H.B. and Wirsel, S.G.** (2008) The yeast signal sequence trap identifies secreted proteins of the hemibiotrophic corn pathogen Colletotrichum graminicola. *Molecular Plant-Microbe Interactions*, **21**, 1325-1336.

**Liu, Y., Boukhelifa, M., Tribble, E. and Bankaitis, V.A.** (2009) Functional studies of the mammalian Sac1 phosphoinositide phosphatase. *Adv Enzyme Regul*, **49**, 75-86.

**Lu, S., Wang, J., Chitsaz, F., Derbyshire, M.K., Geer, R.C., Gonzales, N.R., Gwadz, M., Hurwitz, D.I., Marchler, G.H., Song, J.S., Thanki, N., Yamashita, R.A., Yang, M., Zhang, D., Zheng, C.,**

**Lanczycki, C.J. and Marchler-Bauer, A.** (2020) CDD/SPARCLE: the conserved domain database in 2020. *Nucleic Acids Res*, **48**, D265-D268.

**Mackey, D., Holt III, B.F., Wiig, A. and Dangl, J.L.** (2002) RIN4 interacts with Pseudomonas syringae type III effector molecules and is required for RPM1-mediated resistance in Arabidopsis. *Cell*, **108**, 743-754.

**Mao, Y. and Tan, S.** (2021) Functions and Mechanisms of SAC Phosphoinositide Phosphatases in Plants. *Front Plant Sci*, **12**, 803635.

**Mascher, M.** (2021) Pseudomolecules and annotation of the third version of the reference genome sequence assembly of barley cv. Morex [Morex V3]: e!DAL - Plant Genomics and Phenomics Research Data Repository (PGP), IPK Gatersleben, Seeland OT Gatersleben, Corrensstraße 3, 06466, Germany.

**Mascher, M., Gundlach, H., Himmelbach, A., Beier, S., Twardziok, S.O., Wicker, T., Radchuk, V., Dockter, C., Hedley, P.E., Russell, J., Bayer, M., Ramsay, L., Liu, H., Haberer, G., Zhang, X.Q., Zhang, Q., Barrero, R.A., Li, L., Taudien, S., Groth, M., Felder, M., Hastie, A., Simkova, H., Stankova, H., Vrana, J., Chan, S., Munoz-Amatriain, M., Ounit, R., Wanamaker, S., Bolser, D., Colmsee, C., Schmutzer, T., Aliyeva-Schnorr, L., Grasso, S., Tanskanen, J., Chailyan, A., Sampath, D., Heavens, D., Clissold, L., Cao, S., Chapman, B., Dai, F., Han, Y., Li, H., Li, X., Lin, C., McCooke, J.K., Tan, C., Wang, P., Wang, S., Yin, S., Zhou, G., Poland, J.A., Bellgard, M.I., Borisjuk, L., Houben, A., Dolezel, J., Ayling, S., Lonardi, S., Kersey, P., Langridge, P., Muehlbauer, G.J., Clark, M.D., Caccamo, M., Schulman, A.H., Mayer, K.F.X., Platzer, M., Close, T.J., Scholz, U., Hansson, M., Zhang, G., Braumann, I., Spannagl, M., Li, C., Waugh, R. and Stein, N.** (2017) A chromosome conformation capture ordered sequence of the barley genome. *Nature*, **544**, 427-433.

**Nakagawa, T., Kurose, T., Hino, T., Tanaka, K., Kawamukai, M., Niwa, Y., Toyooka, K., Matsuoka, K., Jinbo, T. and Kimura, T.** (2007) Development of series of gateway binary vectors, pGWBs, for realizing efficient construction of fusion genes for plant transformation. *Journal of bioscience and bioengineering*, **104**, 34-41.

**Pettersen, E.F., Goddard, T.D., Huang, C.C., Meng, E.C., Couch, G.S., Croll, T.I., Morris, J.H. and Ferrin, T.E.** (2021) UCSF ChimeraX: Structure visualization for researchers, educators, and developers. *Protein Sci*, **30**, 70-82.

**Rapazote-Flores, P., Bayer, M., Milne, L., Mayer, C.D., Fuller, J., Guo, W., Hedley, P.E., Morris, J., Halpin, C., Kam, J., McKim, S.M., Zwirek, M., Casao, M.C., Barakate, A., Schreiber, M., Stephen, G., Zhang, R., Brown, J.W.S., Waugh, R. and Simpson, C.G.** (2019) BaRTv1.0: an improved barley reference transcript dataset to determine accurate changes in the barley transcriptome using RNA-seq. *BMC Genomics*, **20**, 968.

**Snyder, J.T., Gershberg, S., Worthylake, D.K., Harden, T.K. and Sondek, J.** (2006) Crystal structure of Rac1 bound to its effector phospholipase C-β2. *Nature structural & molecular biology*, **13**, 1135-1140.

**Vickers, C., Xue, G. and Gresshoff, P.** (2003) A synthetic xylanase as a novel reporter in plants. *Plant Cell Reports*, **22**, 135-140.

**Voinnet, O., Rivas, S., Mestre, P. and Baulcombe, D.** (2003) Corrected: An enhanced transient expression system in plants based on suppression of gene silencing by the p19 protein of tomato bushy stunt virus. *The Plant Journal*, **33**, 949-956.

**Williams, R.L.** (1999) Mammalian phosphoinositide-specific phospholipase C. *Biochimica et Biophysica Acta (BBA)-Molecular and Cell Biology of Lipids*, **1441**, 255-267.

**Zhong, R. and Ye, Z.H.** (2003) The SAC domain-containing protein gene family in Arabidopsis. *Plant Physiol*, **132**, 544-555.
